# Working memory retention by medial entorhinal cortex low-dimensional neural dynamics

**DOI:** 10.64898/2026.08.18.745500

**Authors:** Li Yuan, Jiarui Wang, Xiaohe Li, Wenxi Li, Johnatan Aljadeff, Jill K. Leutgeb, Stefan Leutgeb

**Author notes:** Correspondence (S.L.), (J.K.L.).

## Abstract

The ability to retain information over tens of seconds is essential for carrying out many routine tasks. Recurrently connected circuits are thought to support memory over these timescales, yet how they maintain and represent remembered information remains unclear. We asked whether the medial entorhinal cortex (mEC), where recurrent attractor dynamics generate grid-cell firing patterns and single toroidal attractor manifolds in two-dimensional environments, also supports low-dimensional neural activity patterns during memory retention. During a working-memory task, delay-period mEC dynamics did not show key features of grid firing in simple environments. Instead, broader mEC populations, not limited to grid cells, expressed multiple sequences and recurring activity patterns. Periodically repeating delay-period sequences retained task-relevant information, including past locations and future turn directions. These findings suggest that the mEC network generates low-dimensional dynamics more broadly, with grid-cell activity representing one manifestation and more complex manifold topologies supporting task-related representations during working memory.

## INTRODUCTION

The ability to retain and manipulate information over intermediate time intervals (seconds to ∼1 minute) is critical for executing routine tasks and for guiding everyday decision-making. Central to this process is working memory (WM)—the maintenance and manipulation of information over brief periods to inform future actions. Distinct cortical areas that are associated with different types of WM have been identified ^1, 2^. In studies focusing on short-term retention (seconds) ^3–12^, it is predominantly the prefrontal cortex and its directly connected brain areas, such as the ventral hippocampus (HPC), that are considered the core circuit of a network for spatial WM ^13–24^. However, the canonical circuit has been shown to have a diminishing role in well-practiced tasks ^5, 25^. Furthermore, the persistent and timed firing patterns that have been described in this circuit for intervals of up to ∼10 seconds ^3–12^ may not be suitable for holding information over longer time periods. Additional specialized computations may thus be needed ^5, 25, 26^. Theoretical models suggest that attractor networks support memory retrieval and retention in a broad range of contexts ^27–37^, including spatial navigation. However, the length of time over which these networks can keep memories online is often not specified.

Attractor dynamics are implemented in medial entorhinal cortex (mEC), and mEC is critical for bridging temporal gaps and for spatial WM ^38–41^. Attractor-like low-dimensional dynamics could thus be central to maintaining task-relevant memory over intervals of tens of seconds. In mEC, the activity of grid cells recorded during two-dimensional (2-D) navigation are restricted to an attractor manifold with toroidal topology, and are therefore thought to primarily code for space ^28, 36, 42–46^. Similarly, during stationary running on treadmills and along 1-D paths in a maze, repeating sequential firing patterns of mEC cells emerge that are consistent with trajectories on a torus ^43, 47–49^. While a single torus is well suited to map continuous space, it is inherently limited in its capacity to flexibly retain multiple memories. Computational theories propose potential solutions, including the partitioning of activity into multiple attractor basins or transiently binding cells that participate in the core attractor with other non-attractor populations ^30, 36, 50–60^. For example, in addition to grid cells in simple 2-D navigation tasks, other entorhinal cells could flexibly and transiently be recruited to attractor manifolds and generate more complex structures of coordinated neural activity. This could result in low-dimensional dynamics which combine features of grid cells, such as periodicity, with additional features needed to store multiple memories. However, whether mEC generates structured, memory-relevant neuronal activity patterns over behaviorally meaningful delay periods remains unknown. Moreover, it is unclear how population activity in memory tasks is organized across different internal states that are known to strongly shape mEC dynamics—such as theta-related locomotion versus non-theta immobility. Delayed spatial alternation tasks are hippocampus-dependent and mEC-dependent ^61, 62^ and provide a controlled paradigm to assess the contribution of entorhinal neural dynamics to WM retention over tens of seconds. Intervals in this time range are well within the realm of WM, but too long for persistent or timed firing patterns to remain informative ^8, 26^. To address whether attractor-like dynamics could bridge this gap, we performed Neuropixels recordings from mEC in rats performing a spatial WM task with 10-s or 30-s delay intervals. This design enabled a direct test of whether mEC delay-period activity exhibits attractor-like organization and can support WM retention.

## RESULTS

To examine neural network dynamics during WM retention over intermediate timescales, we trained rats (5 male and 5 female) in a delayed spatial alternation task that included 10-s and 30-s delay periods. During the delay, rats were spatially confined within a central delay zone and either ran continuously on a treadmill or rested with the treadmill off, which enabled us to distinguish delay periods with different brain oscillation patterns—continuous theta during running vs. mixed oscillations while resting. The four task variants (combinations of treadmill on/off and 10-s/30-s delay) were presented in 10-trial blocks in pseudorandomized order, with two repetitions of each of the four variants per session (**Figure 1a**). After training, we chronically implanted Neuropixels 2.0 probes, with the four shanks inserted along the mediolateral extent of mEC (**Figure S1a**). Recordable sites per probe were selected across the four probe shanks to focus on the superficial layers and to maximize yield.

**Figure 1.**
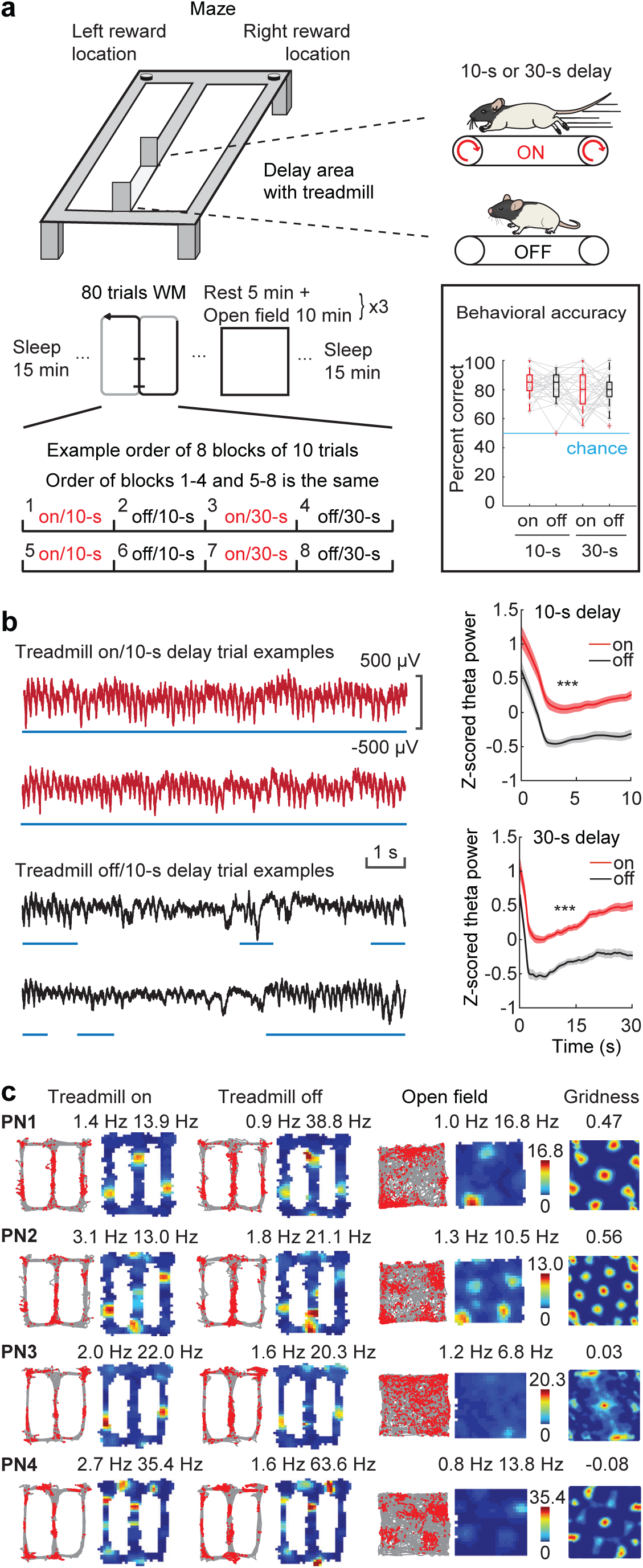
Resting or running on a treadmill during the delay determined the amplitude and persistence of theta oscillations. **a**, Top: schematic of the spatial working memory task. The rat’s behavior during delay intervals was controlled by a treadmill, which was either on or off for a 10-s or 30-s delay period. Bottom: recording session schematic. One recording session included 15-min sleep in the beginning and end, 80 trials in the WM task and three 10-min open fields with 5 min rest in-between. In the WM task, the order of the four treadmill/delay blocks (on/10-s, off/10-s, on/30-s, and off/30-s; 10 trials per block) was shuffled across days, and on the same day, the four blocks were repeated for a second time in the same order. Insert: Spatial working memory task performance in each treadmill/delay condition. No differences were found between conditions (on/10-s: 84.05% ± 1.62, off/10-s: 82.42% ± 1.71, on/30-s: 77.45% ± 2.28, off/30-s: 79.98% ± 1.72; mean ± SEM, n = 33 sessions from 9 rats; F(3,128) = 2.498, p = 0.063, one-way ANOVA). Box plots: central line, edges, whiskers, plus signs indicate median, the 25th/75th percentile, maximum/minimum without outliers, outliers. **b**, Left: Raw LFP traces from one recording site in delay intervals with the treadmill either on (red) or off (black). Blue lines, periods of continuous theta oscillations. Right: Theta power was higher in treadmill-on (red) than in treadmill-off (black) delay periods (z-scored theta power, 10-s delay, on: 0.263 ± 0.056, off: -0.262 ± 0.041; 30-s delay, on: 0.288 ± 0.035, off: -0.291 ± 0.038, mean ± SEM, n = 33 sessions in 9 rats; F(3,128) = 55.62, p = 4.4 x 10^-23^, one-way ANOVA). **c**, Spatial firing patterns of example mEC PNs. Each row shows data from a single neuron. Left two columns: animal trajectories (gray) with spike locations (red) and rate maps for treadmill-on blocks. Numbers above, average and peak firing rates. Middle two columns: the same neuron’s activity during treadmill-off blocks. Right three columns: activity patterns in the open field (trajectory with spike locations, spatial rate maps and spatial autocorrelation map). Numbers above: average rate, peak rate and gridness score. PN, principal neuron. Color bars, firing rate in Hz. *** p < 0.001.

### Recordings from mEC over multiple days

We obtained high-quality recordings over multiple days after surgery, which allowed us to examine neural activity patterns in each rat over 2-5 days with good WM performance (percent correct, on/10-s: 84.05% ± 1.62, off/10-s: 82.42% ± 1.71, on/30-s: 77.45% ± 2.28, off/30-s: 79.98% ± 1.72, mean ± SEM, n = 33 sessions from 9 rats with successful recordings, F(3,128) = 2.50, p = 0.063, ANOVA; **Figure 1a, Table S1**). Most of these sessions (n = 25) were performed with constant treadmill speed, while a subset (n = 8) was performed by switching between two different speeds during treadmill-on trials. Each day, we also recorded the same cells as in the WM task during rest periods and in a random foraging task (3 x 10 min) to classify functional cell types (e.g., grid cells ^42^, spatial non-grid cells ^63, 64^). We began our analyses by examining the rats’ behavior and oscillation patterns. With the treadmill on during the delay, rats continuously ran at the treadmill speed and maintained a stable head position. With the treadmill off, rats were free to move but frequently rested (**Figure S1b** and **c**). As predicted, continuous treadmill running produced sustained theta activity over 10-s and 30-s delays, while the limited movement with the treadmill off resulted in less sustained theta (**Figure 1b, Figure S1d-h**; z-scored power: F(3,128) = 55.62, p= 4.4 x 10^-23^; duration of theta bouts: F(3,128) = 53.72, p = 1.5 x 10^-22^; initial theta bout length after entering the delay area: F(3,128) = 33.35, p = 5.4 x 10^-16^; % time with theta: F(3,128) = 46.63, p = 2.0 x 10^-20^, n = 33 sessions, one-way ANOVA).

### Neural activity patterns on the maze and during the delay interval

To focus on neural mechanisms for WM retention in the task, we recorded activity from 2,642 principal neurons (PNs) in dorsal mEC (31 sessions in 9 rats) and 891 PNs in ventral mEC (13 sessions with and 2 sessions without dorsal mEC, in 5 of the same 9 rats). We first confirmed the expected spatial activity patterns for mEC cells. Most mEC PNs showed stable spatial coding in the WM task and stable spatial firing patterns in the open field, including grid patterns (**Figure 1c, Figure S2**). We next applied UMAP to examine the structure of neuronal activity patterns of simultaneously recorded mEC neurons throughout trials in the WM task. This resulted in low-dimensional manifolds that partially resembled the figure-8 maze, as expected from the spatially stable activity patterns of single cells in the maze. In the same projections, separate state spaces for delay periods with and without continuous running in the delay were observed (**Figure 2, Figure S3**). The distinct neural activity patterns during the treadmill-on delay, treadmill-off delay and the remainder of the maze emerged without major average firing rate differences between conditions (**Figure S4**), which indicates that active sets of mEC neurons were not distinct across phases, but rather exhibited more subtle differences in population activity patterns. When rats were running during the delay, neural activity patterns in UMAP corresponded to extended trajectories that were consistent across trials and connected between the entry and exit from the delay zone. Most progression in UMAP space could be observed in the first 10 s—both in the 10-s delay and the 0-10s interval in the 30-s delay—while it slowed down from 10-30s. Neural activity while resting during the delay displayed a less orderly progression (**Figure 2b**). This suggests that locomotion and associated oscillatory dynamics yielded serial temporal and/or spatial neural activity patterns, as previously reported for entorhinal cells ^39^. To further examine the contribution of locomotion to mEC activity patterns, we compared the firing patterns of single cells between the treadmill-on and treadmill-off conditions.

**Figure 2.**
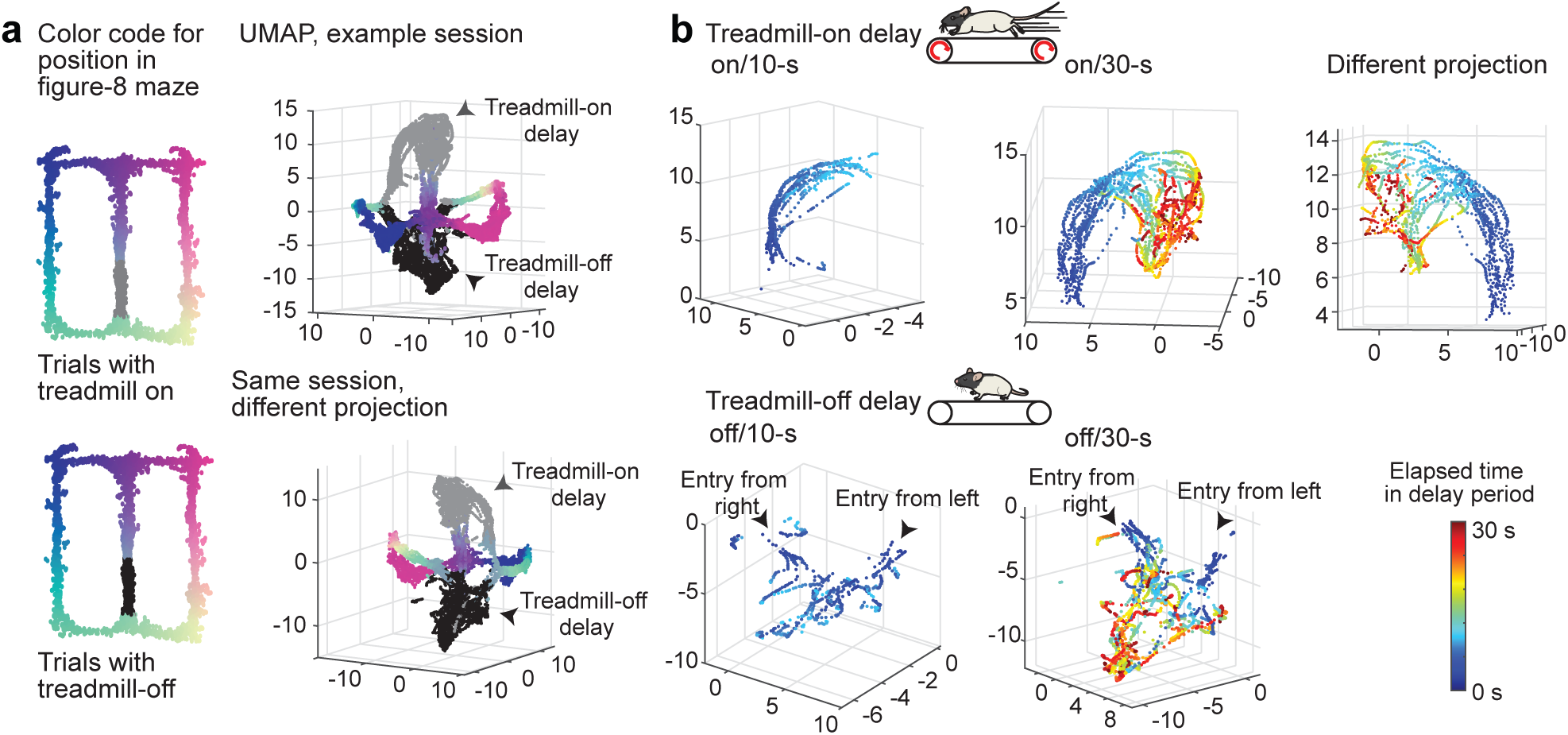
Manifold structure of mEC population activity resembled the maze layout, but with separate regions for treadmill-on and treadmill-off delays. **a**, Left: Schematic of the figure-8 maze with spatial positions color-coded and the delay interval in gray (treadmill-on) or black (treadmill-off). Right: UMAP projection of the activity of simultaneously recorded mEC PNs during the WM task (one session from rat 1146). The two panels on top and bottom show the same 3-D embedding from two different angles, illustrating the separation between maze traversal and delay-related activity. **b**, From the same embedding as in a, the neural trajectories during delay intervals are shown, color-coded by elapsed time within the delay interval. Top, trajectories during treadmill-on delays (left, 10-s delay; middle and right, 30-s delay). Bottom, trajectories during treadmill-off delays (left, 10-s delay; right, 30-s delay).

### Timed activity patterns during delay intervals

To examine the activity of single cells during treadmill-on and treadmill-off delay intervals, we first performed trial-by-trial correlations to determine whether the activity of cells at particular intervals within the delay period depended on running (**Figure 3a**) and sorted cells with consistent activity patterns during the delay by their peak activity (**Figure 3b**). The proportion of consistently active cells showed moderate differences between treadmill-off and treadmill-on trials (**Figure 3c**; dorsal mEC: on/10-s: 33.98% ± 3.09, on/30-s: 36.43% ± 2.66, off/10-s: 27.93% ± 2.27, off/30-s: 30.21% ± 2.19, mean ± SEM, n = 19 sessions; two-way ANOVA: 10-s vs. 30-s: F(1,72) = 0.89, p = 0.35; on vs. off: F(1,72) = 5.98, p = 0.017; interaction: F(1,72) = 0.001, p = 0.97; ventral mEC: on/10-s: 23.45% ± 2.29, on/30-s: 29.36% ± 2.45, off/10-s: 19.07% ± 2.33, off/30-s: 19.63% ± 1.45, mean ± SEM, n = 9 sessions; two-way ANOVA: 10-s vs. 30-s: F(1,32) = 2.50, p = 0.12; on vs. off: F(1,32) = 11.90, p = 0.0016; interaction: F(1,32) = 1.71, p = 0.20), and there were also qualitative differences in activity patterns during the delay. With the treadmill off, peak activity was either early (i.e., within the first 5 s) or cells were persistently active later in the delay interval. During running with the treadmill on, peak times were more evenly distributed, and cells were particularly well ordered during the entire 10-s delay and during the first ∼15 seconds of the 30-s delay (**Figure 3b**). Overall, these findings suggest weaker temporally organized coding during treadmill-off delays while mEC neurons are serially active for at least part of the delay during treadmill-on delays. This aligns with UMAP results showing more consistent trajectories of neural activity with continuous running (see **Figure 2, Figure S3**).

**Figure 3.**
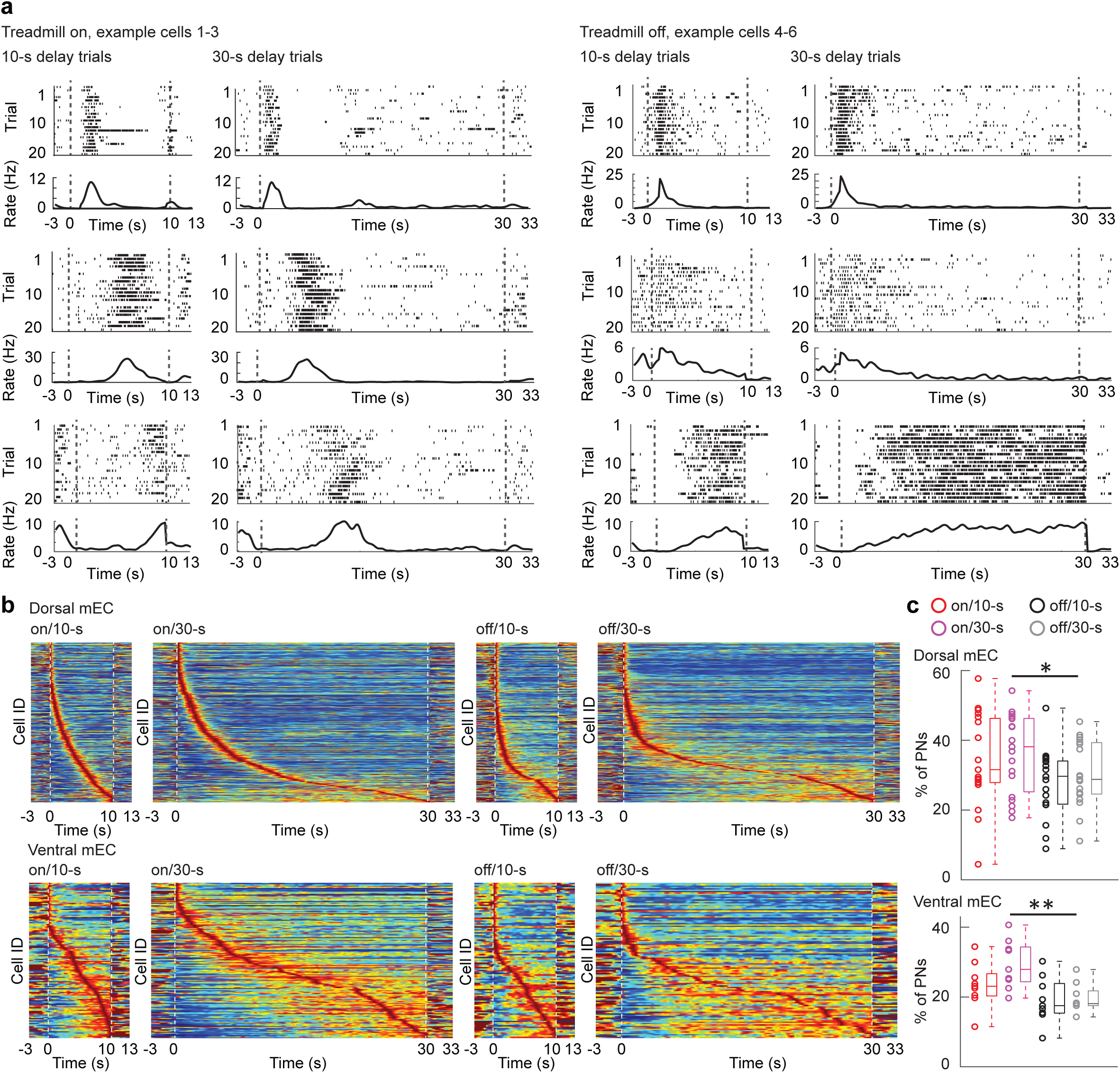
Time/distance coding in mEC across delay conditions. **a**, Example dorsal mEC PNs classified as time/distance cells during treadmill-on and treadmill-off delay intervals. Each panel shows spike rasters (top) and matching firing rate (bottom) for 10-s (left) and 30-s (right) delays. During treadmill-on delay intervals, cells fired at consistent times/distances within the interval, with matching patterns across delay durations. During treadmill-off delay intervals, cells showed time-limited firing only during the first few seconds and were otherwise active throughout the remainder of the delay, often with matching patterns across delay durations. **b**, Activity of the entire population of time/distance cells. Top: dorsal mEC PNs. Of 1968 cells from 23 sessions, 724, 761, 538 and 575 showed consistent firing patterns across trials of the same type (on/10s, on/30s, off/10s and off/30) and are included in the panels. Bottom: ventral mEC PNs. Of 560 cells from 10 sessions, 131, 168, 107 and 112 showed consistent firing patterns across trials of the same type (on/10s, on/30s, off/10s and off/30) and are included in the panels. Cells are sorted by peak firing time within each delay condition. **c,** Treadmill-on delay conditions had a higher fraction of time/distance cells compared to treadmill-off conditions (dorsal mEC: on/10-s: 33.98 ± 3.09%, on/30-s: 36.43 ± 2.66%, off/10-s: 27.93 ± 2.27%, off/30-s: 30.21 ± 2.19%; n = 19 sessions with ≥30 cells from 7 animals; 10-s vs. 30-s: F(1,72) = 0.89, p = 0.349; on vs. off: F(1,72) = 5.98, p = 0.017; two-way ANOVA; ventral mEC: on/10-s: 23.45 ± 2.29%, on/30-s: 29.36 ± 2.45%, off/10-s: 19.07 ± 2.33%, off/30-s: 19.63 ± 1.45%; n = 9 sessions with ≥30 cells from 4 animals; 10-s vs. 30-s: F(1,32) = 2.50, p = 0.124; on vs. off: F(1,32) = 11.90, p = 0.002; two-way ANOVA). * p < 0.05, ** p < 0.01.

To determine whether the ordered activity of mEC neurons preferentially represented elapsed time or distance traveled during treadmill-on delay intervals, we analyzed sessions in which treadmill speed varied across trials. Neurons were classified as either ‘distance cells’ or ‘time cells’ based on whether cross-trial correlations were higher when firing was aligned to distance or time (**Figure S5**). In both dorsal and ventral mEC, there were more distance cells than time cells as measured by significant cross-trial correlations (dorsal mEC: distance cells: on/10-s: 23.93% ± 2.91, on/30-s: 27.72% ± 4.37; time cells: on/10-s: 15.11% ± 2.25, on/30-s: 13.35% ± 3.26, mean ± SEM, n = 8 sessions; two-way ANOVA: 10-s vs. 30-s: F(1,28) = 0.10, p = 0.76; distance vs. time: F(1,28) = 14.35, p = 0.00074; interaction: F(1,28) = 0.84, p = 0.37; ventral mEC: distance cells: on/10-s: 29.07% ± 6.67, on/30-s: 35.75% ± 7.64; time cells: on/10-s: 12.38% ± 1.44, on/30-s: 5.89% ± 1.24, mean ± SEM, n = 5 sessions; two-way ANOVA: 10-s vs. 30-s: F(1,16) = 0.0004, p = 0.98; distance vs. time: F(1,16) = 25.42, p = 0.0001; interaction: F(1,16) = 2.04, p = 0.17). These results indicate that mEC neurons preferentially code for distance during running, which is consistent with the interpretation that mEC neurons path integrate.

### Memory-related firing patterns of single cells during delay intervals

While the finding that mEC cells path integrate in treadmill-on trials is consistent with the interpretation that neural networks are anchored to locations in physical space, it is feasible that firing patterns also depend on past and future trajectories and may therefore retain a memory of task-related variables during the delay. To examine memory-related firing, we quantified the proportion of single neurons with firing rates during the delay that significantly differed when the delay was either part of a left-to-right or a right-to-left trajectory. Firing rate differences were computed in each 5-s bin throughout the delay. Trajectory-selective activity was particularly pronounced early in the delay interval, but its prevalence declined with delay duration, regardless of whether the animal was consistently running or not (**Figure 4a** and **b**; 10-delay: on/10-s: 0-5 s: 11.95%, p = 6.0 x 10^-52^, compared to 5% chance level, binomial test, Holm-Bonferroni corrected, 5-10 s: 6.92%, p = 3.1 x 10^-5^, n = 3105 PNs in 30 sessions with ≤5 errors; off/10-s: 0-5 s: 13.13%, p = 1.2 x 10^-62^, 5-10 s: 9.17%, p = 1.5 x 10^-19^, n = 2834 cells in 28 sessions with ≤5 errors; 30-s delay: see **Table S2**).

**Figure 4.**
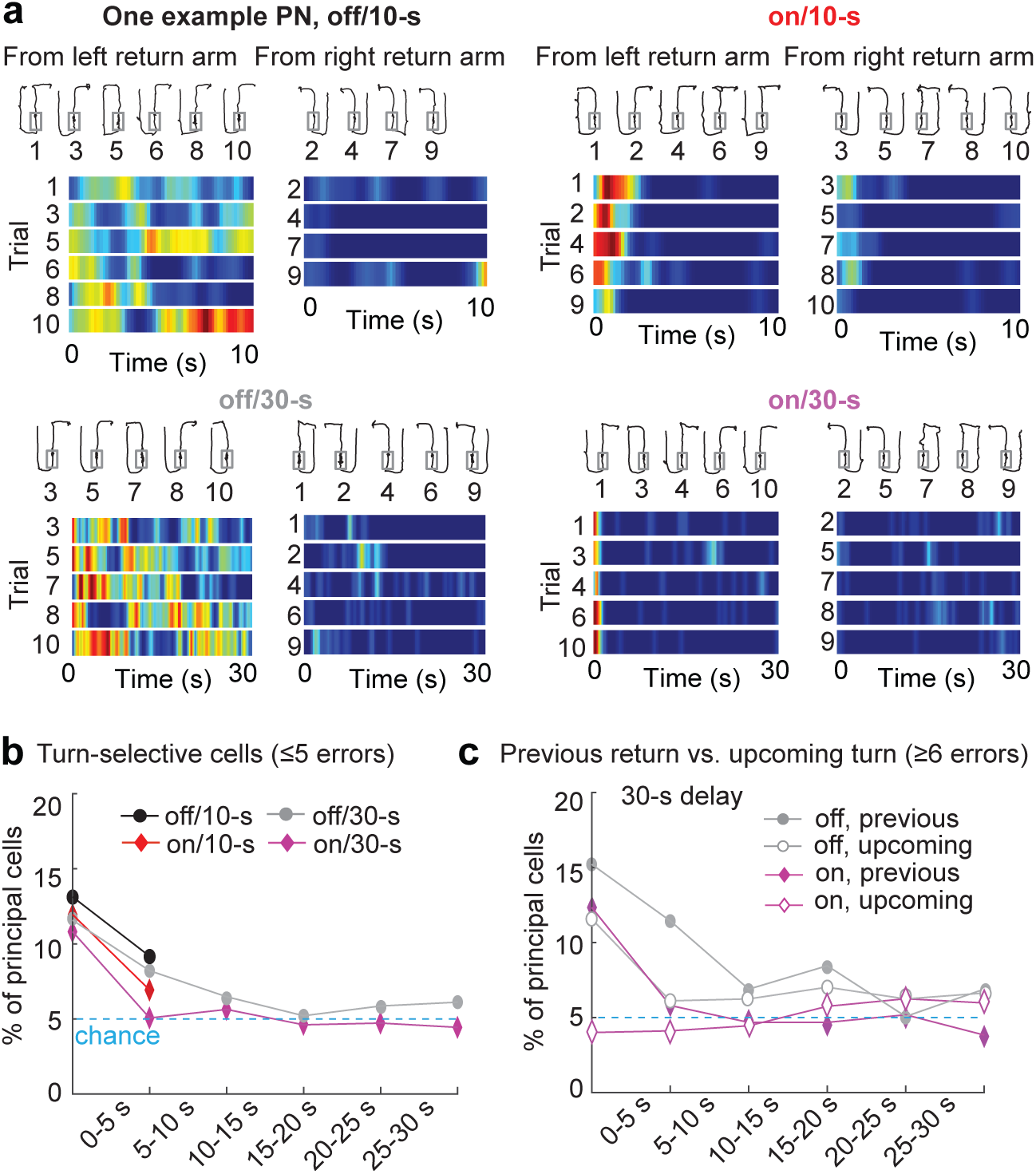
Individual mEC PNs preferentially coded for past return arms. **a**, One example neuron that was more active during delay intervals (off/10-s, on/10-s, off/30-s, and on/30-s) following left-arm returns. Top of each panel: animal trajectory (black line) with the delay area overlaid (gray box). Bottom of each panel: Trial-wise firing rate heatmaps of delay period activity. Color scale: blue to red, 0 Hz to maximum rate. **b**, Trajectory information was retained during the initial portion of the delay in a significant fraction of individual cells (10-delay: on/10-s: 0-5 s: 11.95%, p = 6.0 x 10^-52^, compared to 5% chance level, binomial test, Holm-Bonferroni corrected, 5-10 s: 6.92%, p = 3.1 x 10^-5^, n = 3105 PNs in 30 sessions with ≤5 errors; off/10-s: 0-5 s: 13.13%, p = 1.2 x 10^-62^, 5-10 s: 9.17%, p = 1.5 x 10^-19^, n = 2834 cells in 28 sessions with ≤5 errors; 30-s delay: see **Table S2**). **c**, In 30-s delay intervals with the treadmill on, mEC PNs were selective for the previous return arm, but not for the upcoming turn direction (see **Table S2** for statistics).

To examine whether memory-related delay-period activity reflected previous maze arm location (‘retrospective’) or upcoming turn direction (‘prospective’) coding, we selected session blocks with a large number of error trials (≥6 errors in 20 trials, n = 15 sessions for the on/30-s condition and 6 sessions for the off/30-s condition) in which these cases could be dissociated. A larger proportion of mEC neurons coded for previous location than for the upcoming turn. In particular, prospective coding was not observed in the treadmill-on condition (**Figure 4c**, see **Table S2** for statistics). Because memory coding in single cells was limited, in particular while running on the treadmill during the delay, we next focused on population-level organization of mEC neural activity patterns during running. In particular, we asked whether population dynamics by mEC PNs code for memories more robustly than single cells.

### Dorsal mEC cells were periodically active during running throughout the delay interval

A prominent feature of grid cell attractor manifolds is their periodicity in space. Yet, our initial analysis that aligned mEC single-neuron activity to treadmill distance did not reveal periodicity. A possible reason for trial-averaged analyses to not show periodicity are trial-to-trial inconsistencies in the onset of periodic firing, as evident for a subset of example mEC cells (**Figure 5a** and **b**). Therefore, we next used spatial autocorrelations—using the data from the 30-s delay intervals—to determine whether mEC neurons exhibited periodic firing patterns above chance levels. In dorsal mEC, 24.26% ± 2.20 (n = 27 sessions, mean ± SEM) neurons showed significant periodic activity, while in ventral mEC periodic firing was less prominent (**Figure 5c**; 11.75% ± 2.32, n = 14 sessions, p = 0.0007; two-sample t-test). For dorsal mEC neurons, periodic distance coding during the delay and grid firing in the open-field occurred in partially overlapping cell populations (35.1% periodic cells are grid cells. 37.5% grid cells are periodic cells; **Figure 5d**). For cells exhibiting both properties, the periodic distance was positively correlated with the grid spacing of that cell (**Figure 5e**; r = 0.29, p = 8.1 x 10^-6^, n = 232, Spearman’s correlation), although absolute periodic distance and grid spacing substantially differed.

**Figure 5.**
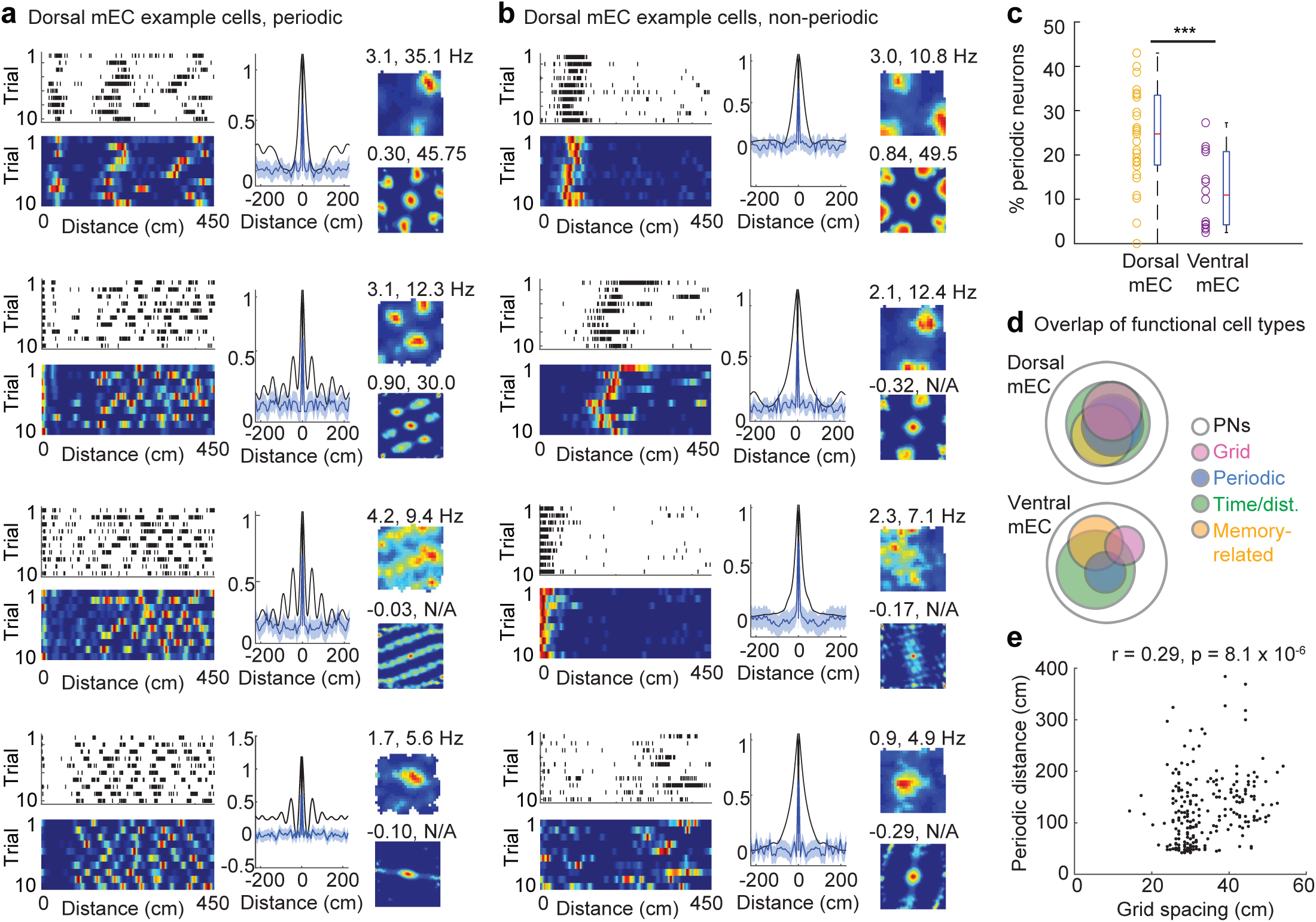
Periodic firing of mEC PNs during running on the treadmill in the 30-s delay interval. **a**, Four example dorsal mEC PNs that were periodically active. Left top and bottom of each panel: spike rasters and color-coded distance tuning curves. Middle: autocorrelogram of the distance rate function (black) compared with an autocorrelogram from shuffled data (blue line, mean; blue shading, 99% confidence interval). Right: open-field rate map (color scale: blue to red, 0 Hz to maximum rate, with mean and peak firing rates on top) and spatial autocorrelation map (with grid score and spacing on top; spacing is N/A for non-grid cells) of the same cell. **b**, Examples of non-periodic dorsal mEC PNs, displayed as in a. **c**, The proportion of periodic cells was significantly higher in dorsal mEC (24.26% ± 2.20 of 2587 PNs, n = 27 sessions from 8 animals) than in ventral mEC (11.75% ± 2.32 of 863 PNs, n = 14 sessions from 5 animals; two-sample t-test, p = 0.0007). **d**, Overlap of functional cell types in dorsal mEC (top) and ventral mEC (bottom). Significant overlap is observed between time/distance cells (see Figure 3b), memory-related cells (see Figure 4b), periodic cells and grid cells in dorsal mEC (adjusted permutation test, p < 0.05 for all comparisons, see **Figure S11d** for detailed statistics). **e**, Periodic distance (measured during running in the 30-s delay) and grid spacing (measured in the open field) were correlated (Spearman’s r = 0.29, p = 8.1 × 10⁻⁶, n = 232 neurons). *** p < 0.001.

### MEC neurons organized into repeating sequences during delay intervals

Given that a proportion of single mEC cells was periodically active during running in the delay area without visual cue change, we reasoned that the entire population of mEC cells may organize into periodically recurring sequences during treadmill running, as previously reported for running along one-dimensional trajectories and on treadmills in darkness ^47, 49, 65^. To investigate whether mEC PNs organized into sequentially structured activity patterns, we applied seqNMF to neural activity data during delay intervals (**Figure 6a** and **b, Figure S7**). This analysis revealed 93 distinct sequences in 24 sessions from 8 rats, and each of these sequences typically repeated multiple times throughout 30-s delay intervals (**Figure 6c-f, Figures S8, S9, S10** for additional examples from three different rats). The large number of sequences already indicates that sequences were not as stereotyped as predicted by a purely spatial account for sequence generation, and properties of sequences were thus analyzed in more detail.

**Figure 6.**
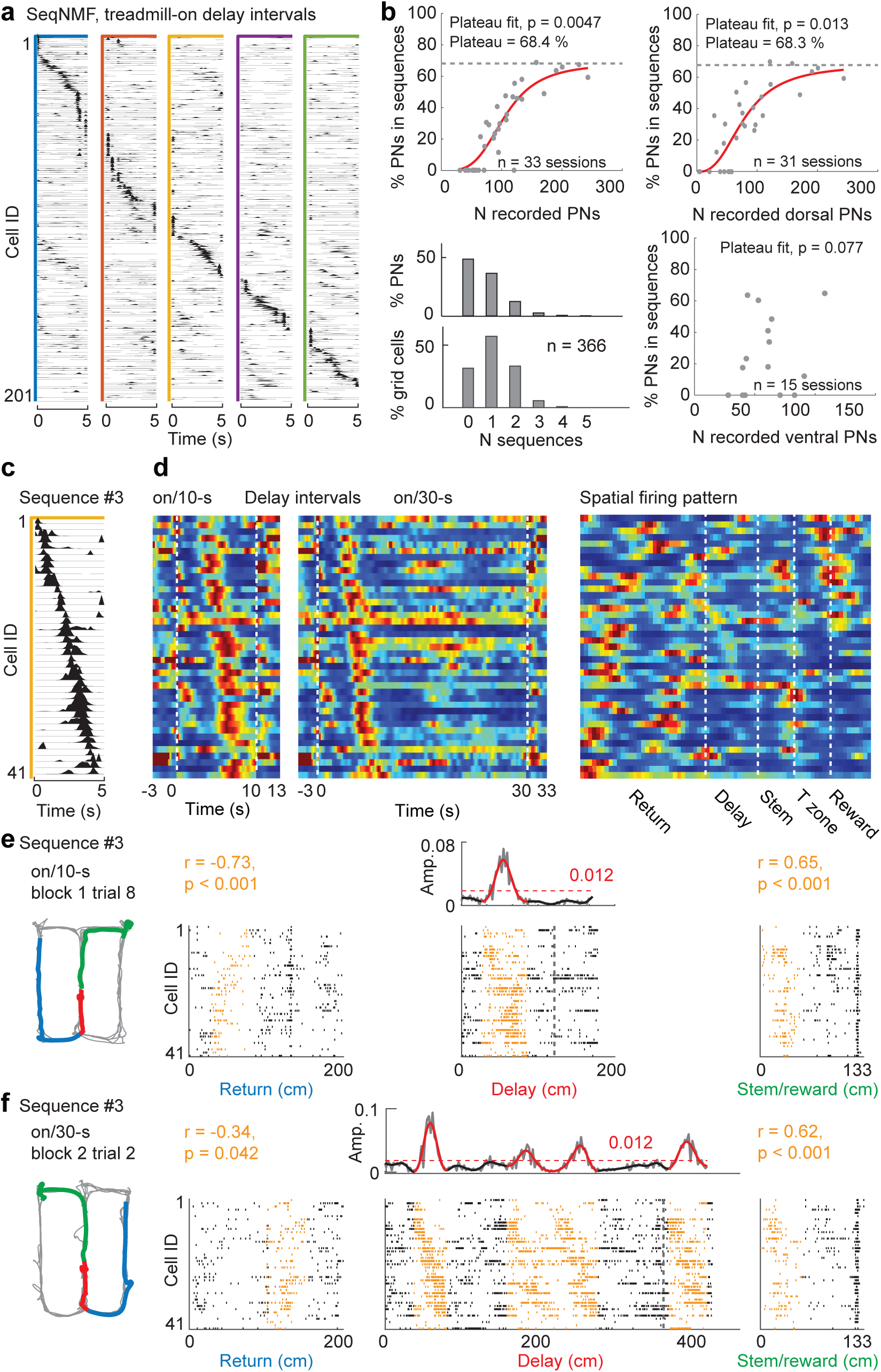
MEC PNs organized into repeating sequences during delay intervals. **a**, Sequences were detected by applying seqNMF to mEC PN activity during treadmill-on delay intervals. Each of five detected sequences is framed with a different color, and PNs within sequences are ordered by the timing of their peak weight. **b**, Top left: Relationship between the total number of recorded mEC PNs and the fraction participating in at least one sequence. PN participation in sequences approached a plateau at ∼68% (non-linear saturation fitting, p = 0.005). Top right: Dorsal mEC PN participation in sequences, also reaching a plateau at ∼68% (non-linear saturation fitting, p = 0.013). Bottom right: Ventral mEC PN participation in sequences, which did not reach a significant plateau (non-linear saturation fitting, p = 0.077). Bottom left: Distribution of the number of sequences in which mEC PNs and grid cells participate. Most PNs contributed to zero or one sequence, whereas smaller fractions participated in multiple sequences. Note that the fractions here include all sessions, irrespective of whether PN participation in sequences was close to the plateau. **c**, PNs participating in the third sequence from the example session shown in a, after selecting neurons with weights above chance (see **Figure S7**). **d**, Average temporal and spatial firing patterns (left: on/10-s; middle: on/30-s, right: linearized figure-8 maze) of neurons from the example sequence, ordered as in c. **e**, Occurrence of the example sequence in c during a trial with 10-s delay. From left to right: animal’s trajectory color-coded by maze segment (blue, return; red, delay interval; green, stem to reward), sequence occurrence in reverse order during return (rank-order correlation value and significance on top), sequence occurrence in the delay interval (Top: gray line, sequence amplitude; black line, smoothed amplitude, turns red throughout detected events; red dashed line and number, chance level; bottom: spike rasters for each neuron in the sequence, ordered as in c; vertical dashed line, opening of barrier at end of delay time), sequence occurrence in forward order in stem to reward (rank-order correlation value and significance on top). **f**, Repeated sequence occurrences throughout the delay in an example trial with 30-s delay, depicted as in e.

The number of sequences, the number of PNs per sequence, and the number of PNs in any sequence all scaled with the total number of mEC PNs recorded per session (**Figure S11a-c**). The fraction of mEC PNs, and in particular of dorsal mEC PNs, participating in any sequence plateaued at ∼70%, suggesting a saturation point in recruitment (**Figure 6b**). In contrast, ventral mEC neuron participation metrics did not scale with the number of recorded ventral PNs, suggesting that sequence features are primarily governed by dorsal mEC dynamics. For mEC PNs included in sequences, the majority participated in only one sequence and increasingly smaller proportions participated in two or more. The same pattern was seen for grid cells, but overall, grid cells had a higher likelihood of participating in sequences than other PNs such that many grid cells participated in two or more sequences (**Figure 6b**). Together, these results suggest that dorsal mEC strongly contributes to the emergence of structured, repeating sequences during working memory delays. Interestingly, dorsal mEC neurons participating in sequences are a much larger proportion of neurons than cells in other categories, including grid cells (**Figure S11d**), which indicates that attractor-like dynamics in memory tasks extend beyond the sub-network that is participating in grid cell attractors.

### Independent sequence dynamics during the delay period

Multiple distinct neuronal sequences were detected within the same delay intervals (**Figure 7a, Figure S11a** and **S12**). It is feasible that the multiplicity of sequences during running in delay periods emerged from fragmentation of a single sequence by the seqNMF method. If this were the case, the fragments should have a consistent temporal relation to each other. Examination of temporal relationships revealed that most sequence pairs did not exhibit consistently timed correlations (**Figure 7b, Figure S12**), indicating that each sequence operated largely independently rather than as part of a coordinated sequence-of-sequences. Despite the variability of individual sequence occurrences, the likelihood of observing a sequence event and the average quality (i.e. amplitude) of a sequence remained remarkably stable across time (**Figure 7c**). Thus, the overall level of sequence activity did not decline across the delay period. This contrasts with the gradual reduction in the occurrence of single-cell time/distance coding (see **Figure 3**) and suggests that population-level sequence dynamics are maintained across the WM delay intervals, even when individual neurons cease to show patterns that are aligned to time or distance.

**Figure 7.**
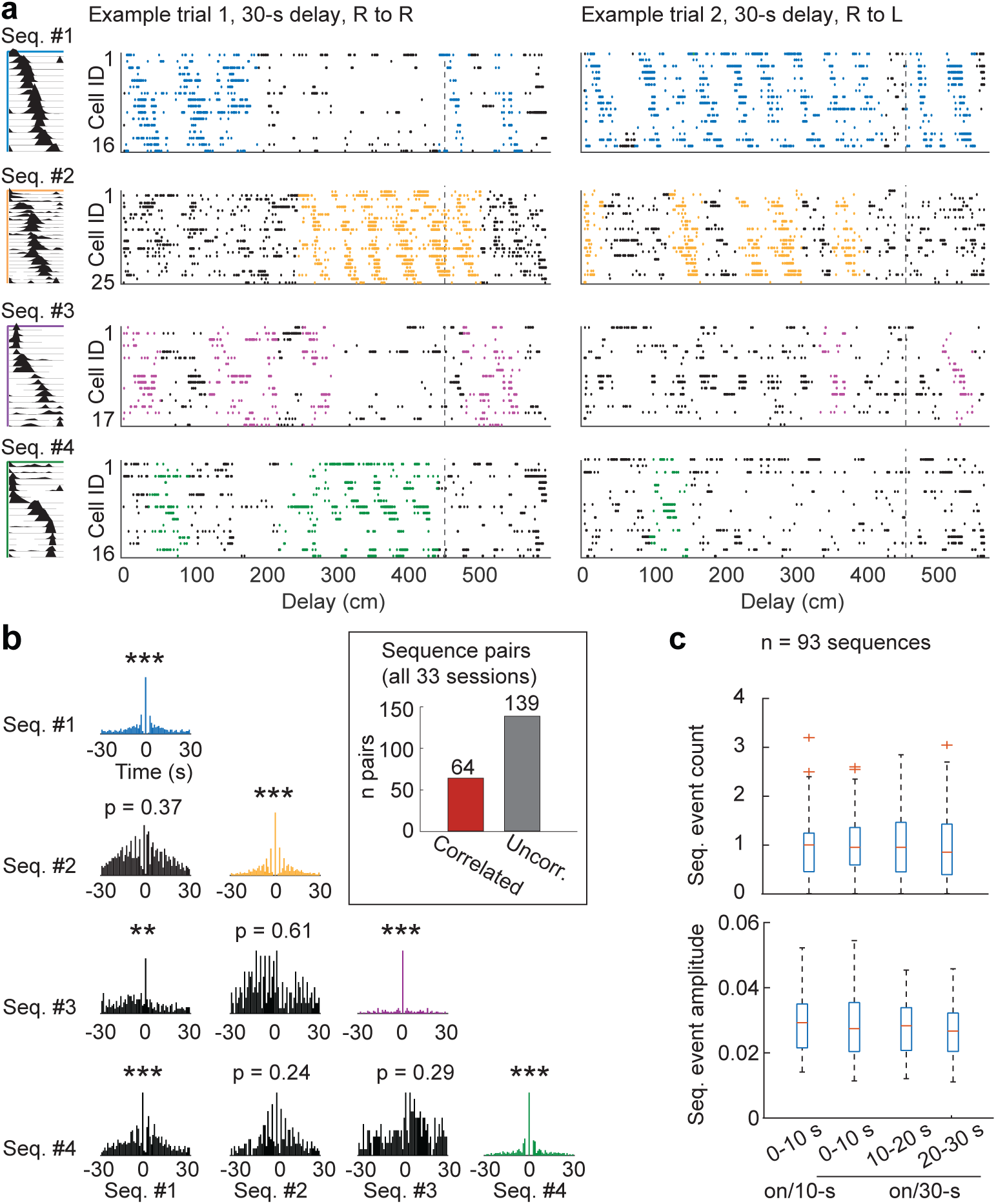
Multiple distinct mEC sequences repeated throughout delay intervals. **a**, Two example trials with four distinct sequences throughout the delay intervals. For each sequence, the left panel shows the SeqNMF-derived weight profile across PNs, and the panels to the right show spike rasters of participating neurons, ordered as shown to the left. **b**, Autocorrelations reveal robust temporal structure within individual sequences, whereas cross-correlations show only limited temporal coupling between different sequences. Insert: Across all sequence pairs, only a subset (64 of 203) exhibited significant temporal correlations, indicating that the majority of sequences were temporally independent. **c**, Top: Average sequence event counts remained consistent across 10-s segments in delay intervals [10-s delay: 0.96 ± 0.07; 30-s delay, 0-10 s: 1.00 ± 0.06; 10-20 s: 1.02 ± 0.07; 20-30 s: 1.00 ± 0.08, n = 93 sequences; one-way ANOVA, F(3,368) = 0.109, p = 0.955]. Bottom: Peak amplitudes of sequence events remained consistent across 10-s segments in delay intervals [10-s delay: 0.029 ± 0.001; 30-s delay, 0-10 s: 0.028 ± 0.001; 10-20 s: 0.028 ± 0.001; 20-30 s: 0.027 ± 0.001, n = 93 sequences; one-way ANOVA, F(3,357) = 0.913, p = 0.435]. ** p < 0.05, *** p < 0.001.

### MEC sequences spanned the entire WM task

To test whether the sequences that were detected during treadmill running also occurred during other task phases, we compared the order of neurons in detected sequences to their activation patterns during treadmill-off delays and during traversal of the return arm (preceding the delay) and the stem-to-reward segment (following the delay; **Figure 6c-f, Figure S13a**). Corresponding sequences to those that occurred during running in the delay were also generated while running in stem-to-reward and during treadmill-off delays. Sequences in the stem-to-reward unfolded over a similar time scale to those in the treadmill-on delay. In contrast, sequences generated during the treadmill-off delays were temporally compressed from ∼5 s to hundreds of milliseconds (**Figure S13d**). Given that entorhinal sequences could show the time-compression that is well-established for ripple-related hippocampal firing, we also tested an additional feature of ripple-related activity, which is their occurrence in reverse order. Notably, the temporal order of activation was reversed when sequences played out during running on the return arm (**Figure 6c-f; Figure S9, S10** and **S13**). While this resembles a feature of hippocampal activity, the reverse order of entorhinal activity did not occur in a time-compressed manner, but rather on a time scale that was approximately matched to the activity during treadmill running. To confirm that the correspondence in activation patterns across task phases was robust, we quantified the correspondence to treadmill-on sequences for each of the different types of activity patterns. Of the sequences detected during treadmill-on periods, 95.7% recurred in reverse order on the return arm at least once, 94.6% in the original order in stem-to-reward and 51.6% as time-compressed replay during treadmill-off delays (**Figure S13c**). Together, these results demonstrate that mEC delay-period sequences span the entire working-memory task structure. They reflect recent past trajectories, predict upcoming future paths, and persist—albeit in compressed temporal form—during delays without running. This suggests that mEC sequences provide a continuous population-level representation linking past and future behavior across phases of the working-memory task.

### Sequential firing during delay intervals was memory-related

We next examined whether the complex pattern consisting of multiple sequences generated in mEC during treadmill-on delay intervals represented a memory of task-related variables. Specifically, we quantified whether sequence occurrence above a threshold amplitude from shuffled data selectivity depended on the preceding return arm, the upcoming turn direction or the accuracy of the impending choice (**Figure 8a** and **b**). These analyses were performed in each of the 10-s segments of the 30-s delay (**Figure 8c, Figure S14**). In the initial 0-10 s period, occurrence of a subset of sequences depended on whether the rat came from the left versus right return arm. During the middle 10-20 s segment, sequences selectively occurred depending on the upcoming choice direction. In the final 20-30 s segment, sequence amplitude reliably differentiated correct from error trials. Thus, mEC population sequences differ in representational content over the course of the delay—from coding for past trajectories, to predicting future choices, and ultimately, to signaling the likelihood of correct performance. Low-dimensional mEC dynamics are therefore not strictly governed by space, but also include memory-related content and a representation of future choice.

**Figure 8.**
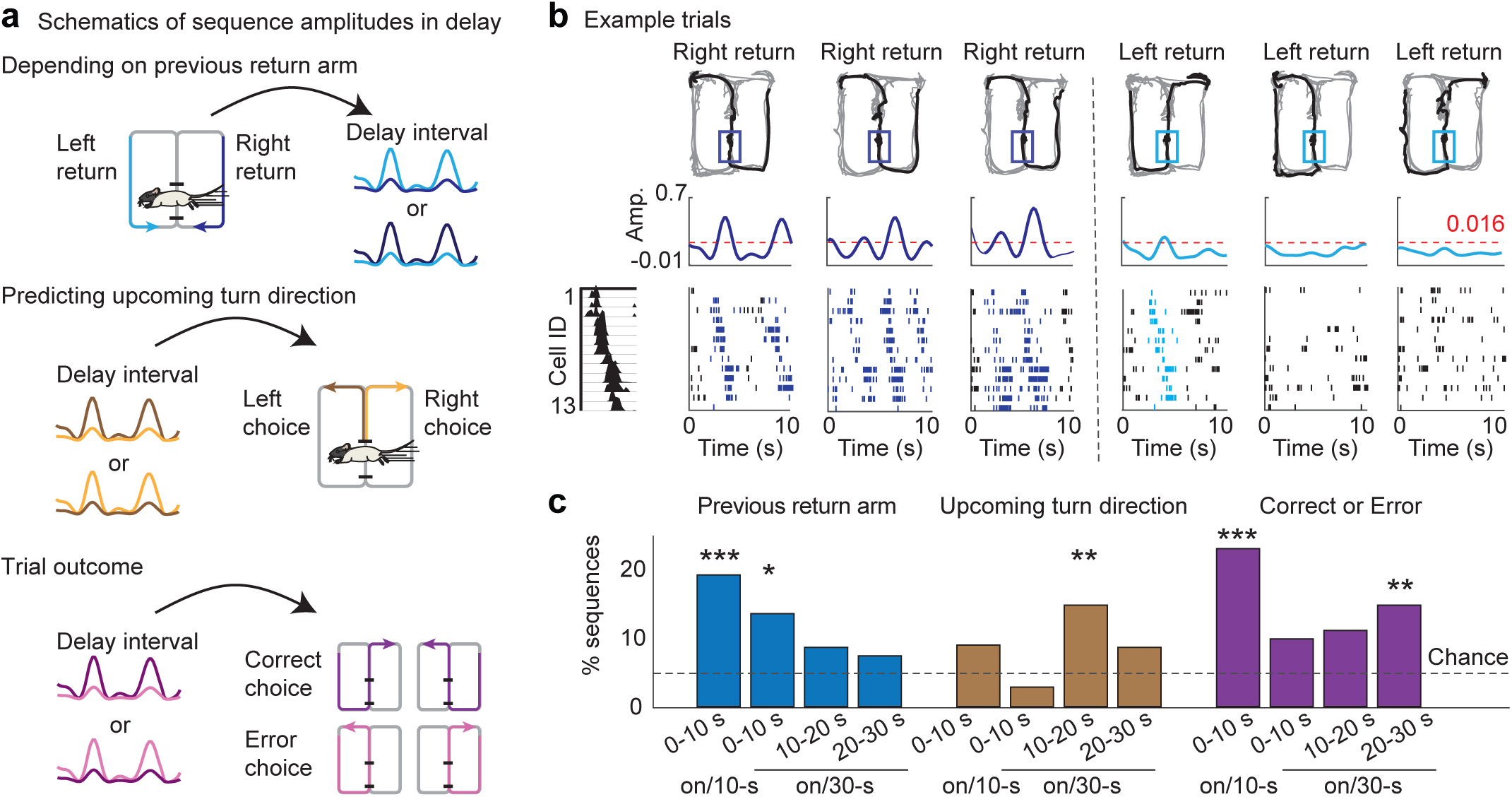
Delay-period sequences flexibly coded for memory-related information. **a**, Schematics illustrating that treadmill-on delay sequences can preferentially be active during delay intervals depending on the preceding return arm (left vs. right), the upcoming turn (left vs. right), or the trial outcome (correct vs. error). **b**, Example sequence with return arm-dependent activity (measured as amplitude modulation). Top: Behavior trajectory (gray, all trials; black, single trial) with delay zone highlighted (dark and light blue when coming from either the right or left return arm). Middle: sequence amplitude during the delay interval. Bottom: spike rasters of contributing neurons, ordered as shown in the sequence template to the left. **c**, Different types of memory-related information were expressed across the delay period (Return left vs. right: on/10-s: 19.23%, p = 0.00007, n = 78; on/30-s: 0-10 s, 13.58%, p = 0.019; 10-20 s, 8.64%, p = 0.51; 20-30 s, 7.41%, p = 0.60, n = 81; upcoming turn left vs. right: on/10-s: 8.97%, p = 0.57, n = 78; on/30-s: 0-10 s, 3.74%, p = 0.80; 10-20 s, 14.82%, p = 0.007; 20-30 s, 8.64%, p = 0.38, n = 81; trial outcome: on/10-s: 23.08%, p = 5 x 10^-7^, n = 78; on/30-s: 0-10 s, 9.88%, p = 0.39; 10-20 s, 11.11%, p = 0.14; 20-30 s, 14.83%, p = 0.006, n = 81; binomial tests compared to chance level of 5%, p values corrected, trial blocks with ≥3 errors in 20 trials included). * p < 0.05, ** p < 0.01, *** p < 0.001.

## DISCUSSION

The use and processing of items in WM is essential for performing routine tasks and for everyday decision making. To test whether attractor networks in mEC could contribute to WM retention over intermediate intervals, we performed multi-shank Neuropixels recordings to monitor large ensembles of mEC neurons as rats performed a spatial alternation task requiring WM over 10-s or 30-s delay intervals. By implementing two distinct behavioral conditions during the delay—enforced locomotion on a treadmill and voluntary immobility—we could directly test how mEC delay-period dynamics depended on running behavior and associated network oscillations. Activity of mEC neuron populations formed neural manifolds that encompassed the spatial structure of the task and the two delay conditions. Notably, the treadmill-on and treadmill-off delay intervals were represented as separate from activity in the remainder of the maze. Furthermore, the two types of delay intervals were distinct from each other, indicating a diversity in coding task-relevant information. We then analyzed the delay interval in detail to examine the organization of single-cell and population activity patterns. When rats were running on a treadmill during the delay interval, individual mEC cells coded for fixed or periodic distances, while memory-related coding was prominent in only the first few seconds of the delay interval. At the population level, mEC delay-period activity exhibited repeating sequences. However, unlike random foraging in 2-D environments, where the activity of a subpopulation of mEC cells maps onto a single toroidal attractor ^43^, more complex low-dimensional network dynamics were observed during working memory retention. Multiple sequences formed, which together included a much larger mEC population than just the grid cell population. Each of the sequences recurred throughout the delay interval, and the strength of manifestations of sequences during the delay interval could code for memory-related information, such as past locations and future turn direction. The activation of sequential neural activity, possibly along an attractor manifold, can thus support WM retention across intermediate timescales.

Many theoretical models have proposed that attractor dynamics are useful for memory retrieval and for holding information in memory. Attractors can manifest as various neuronal activity patterns ^32^. First, activity patterns can converge and remain at a local or global minimum. Such point attractors are characterized by at least briefly displaying stationary activity. In the WM task, persistent activity may be a manifestation of this type of attractor, which is typically observed in studies where subjects were mostly stationary during the delay. Correspondingly, we observed many cells during the treadmill-off delay intervals which could be characterized as persistently firing. When networks display this type of activity, even the analysis of single cells that were participating in this scheme, revealed memory-related activity. Second, attractor surfaces can correspond to low dimensional manifolds, and one example are toroidal manifolds that emerge in populations of mEC cells that map 2-D space. Moving along variable trajectories on a toroidal manifold results in grid cells ^42, 43^ and taking stereotyped trajectories—either on the manifold or in real space—results in repeating activity patterns. Accordingly, toroidal manifolds can be interpreted to include a memory component—integrating the path that has been taken in an environment ^28^. However, such an attractor network is in its simplest form rigidly tied to space and too inflexible to store even a limited number of distinct items. These types of networks nonetheless have the desirable property that they can maintain information for extended time periods, if they could be extended to allow for the repeating activity patterns to correspond to more than 2-D space.

In our recordings during delay intervals, we found that single-cell mEC activity showed periodic activity with distance, but analyses of population activity revealed far more complex patterns. There were multiple repeating sequences that were not related to each other in a stereotypic way, which rules out the possibility that our analysis merely fragmented a single sequence into multiple segments. Importantly, the fraction of mEC cells participating in the sequence was much higher than the fraction of identified grid cells in our study and was even higher than the fraction of grid cells reported with highly inclusive methods of grid cell identification ^66^. This suggests that many cells in the mEC network can participate in generating repetitive firing patterns and that grid cells form a subnetwork within a larger circuit which can generate these dynamics. It is of course possible that grid cells are the cells that make up a core attractor—consisting of the most strongly connected recurrent cells—and that other mEC cells can be flexibly tied to the core to generate a larger network with attractor-like dynamics that retains non-spatial task-relevant information. However, these dynamics would imply that—during memory retention in the WM task—grid cells do not remain inherently coherent with each other, but partially dissociate to contribute to one of the sequences. Irrespective of the contribution of grid cells to each of the sequences, emerging network dynamics during memory retention would only be consistent with the mEC activity patterns forming a single torus, if virtual paths on the manifold are highly variable and not directly determined by space. Alternatively, the surface of the manifold may assume a more complex shape than a torus ^36, 67^, such that the more complex low-dimensional neural activity only partially maps to 2-D space and is instead better compatible with multiple memory states.

Importantly, we show here that multiple repeating sequences can be generated flexibly. We observe variability of sequence manifestation across trials and that individual sequences can be more strongly expressed in the beginning or towards the end of the delay. Sequences nonetheless occur in a balanced manner across the delay interval, such that—on average—the likelihood of observing a sequence was approximately equal throughout. This is in contrast to findings for single hippocampal cells, which can be predominantly observed to be well-timed during only the beginning of the delay intervals^26, 68^. In parallel, individual mEC cells also predominantly show memory-related coding in only the beginning of the delay, and this effect was particularly pronounced when animals were running during the delay. In contrast, sequences could recur throughout the delay interval and carried task-relevant information about both past and future behavior. Early in the delay, sequence amplitude reflected the previous return-arm direction, while later in the delay, it predicted the upcoming choice direction and the correctness of the forthcoming choice. Thus, mEC dynamics flexibly represent different content over time, including retrospective, prospective and performance-related signals. The importance of sequential network organization is apparent in the finding that standard single cell analysis methods did not reflect these network dynamics.

Taken together, the memory-related dynamics of mEC network activity reveal that a large fraction of the mEC network is fundamentally designed to generate low-dimensional neural dynamics. Manifestation of mEC dynamics as a toroidal manifold that gives rise to grid cells may only be one of many subspaces that these networks can implement. In more complex tasks, the repetitive activity features of an attractor network are retained, but may assume more complex manifold shapes that accommodate task-related representations other than being strictly tied to 2-D space. These findings reconciliate the unequivocal role of mEC for inflexibly representing 2-D space with the importance of entorhinal cortex for memory processing. While both cognitive functions can be supported with low-dimensional neural dynamics, a larger network than just the grid-cell attractor contributes to memory-related functions.

## ACKNOWLEDGEMENTS

We thank Mia Anderson and Deborah Chen for technical assistance. This work was supported by NIH grants NS086947, NS102915, NS121231 to S.L., NSF grant 2024776 to S.L. and NIH grant MH119179 to J.K.L.

## AUTHOR CONTRIBUTIONS

S.L., J.K.L., and L.Y. designed the experiments, conceptualized analyses, interpreted data and wrote the manuscript. L.Y. developed the recording methods, acquired and processed data, designed analyses, wrote and curated analysis code and analyzed the data. J.W., X.L. and W.L. acquired and processed data, J.A. designed analyses and edited the manuscript. S.L. and J.K.L. supervised the project.

## CODE AVAILABILITY

Custom code for processing the data reported in this paper will be available from Github. Standard code that is not novel to this study, if not already publicly available, will be available from the corresponding authors upon request.

## DATA AVAILABILITY

Source data will be provided with this paper. Additional data that support the findings of this study are not in standardized formats, but will be made available without restrictions upon request to the corresponding authors.

## METHODS

### Approvals

All experimental procedures were conducted at the University of California, San Diego according to National Institutes of Health guidelines and were approved by the Institutional Animal Care and Use Committee.

### Animal subjects and behavioral task

Experiments were conducted in 10 Long-Evans rats (5 male, 5 female), housed individually under a reverse 12 h light/dark cycle. During behavioral training and recording phases, rats were food restricted to ∼85% of the *ad libitum* weight. The behavioral apparatus was a white figure-8 shaped maze (150 cm x 100 cm) constructed from white acrylic sheets. All sections of the figure-8 maze were composed of 10-cm wide tracks with 2-cm tall ridges on each side. The delay area (70 cm long, 10 cm wide) was located in the half of the center arm that the rat first entered. A treadmill (Harvard apparatus) was fitted into the delay area as the floor for the rats to rest or run on (see **Figure 1**). During the delay periods, the rats were spatially restrained in the delay area by an automatic barrier at the front of the delay zone and by a manual barrier behind its tail. The manual barrier was placed behind the rats after they entered the delay area.

We trained 6-10 weeks old rats to collect chocolate milk at two corners of the figure-8 maze (i.e., the reward locations), with food only available at the reward location if the rats chose the opposite turn in the T-zone compared to the previous trial. During pretraining, rats were given 5 days to habituate and explore the maze freely. Next, they were trained to perform a continuous version of the alternation task for 2-4 weeks until they reached ≥90% accuracy in two of three consecutive training days. Then, the delayed version of the task with and without treadmill running during the delay was introduced to all rats, and their training continued for 10 days before electrode implantation. Rats were trained with four trial types by combining two delay interval durations (10 s or 30 s) with the two treadmill states (off or on) during the delay. Ten consecutive trials with one of the four trial types were conducted before switching to a different trial type. The order of trial types was shuffled across training/recording days, and within each day, the four blocks were repeated for a second time in the same order (**Figure 1**). After completing the working memory task, rats were exposed to three cycles of open field exploration (10 min, 1.2 m × 1.2 m box) separated by 5-min rest periods. In addition, each recording session began and ended with a 15-min sleep period.

In a subset of recording sessions, all 10 consecutive trials under the treadmill-on delay condition were conducted at a constant speed, ranging from 12 to 15 cm/s. In other sessions, the treadmill speed differed between the first and second sets of 5 trials—using combinations of 10 cm/s and 20 cm/s or of 15 cm/s and 20 cm/s. The order of the low-speed and high-speed trials was randomly varied across sessions. The speed variations were implemented to dissociate the effects of elapsed time from distance traveled during the delay period.

### Surgery and recording methods

Two Neuropixels 2.0 probes (IMEC) were chronically implanted in mEC and the medial prefrontal cortex of the same animal with retractable metal drives (R2Drive, 3Dneuron). A total of 10 rats underwent this procedure. Anesthesia was induced with 2-3% isoflurane (in oxygen at 1 L/min), and analgesia was provided with buprenorphine (1 mg/kg, subcutaneous). After placing skull screws—one with a silver ground wire (AGW1010, World Precision Instruments) attached to serve as the reference—and performing targeted craniotomies (MEC: 4.6 mm right of the midline; mPFC: 2.0-2.2 mm anterior to bregma, 0.6 mm lateral to the midline), the Neuropixels probes were inserted and secured with dental cement.

Rats were given 4-5 days to recover in their home cages with unlimited access to food and water. Following recovery, they were food restricted and resumed daily sessions in the delayed spatial alternation task. Electrophysiological data were collected using SpikeGLX (version 20230905) through an IMEC PXIe acquisition module installed in a National Instruments PXIe-1071 chassis (gain: 100; sampling frequency: 30,000 Hz). Position tracking data were recorded using a Digital Lynx system (Neuralynx, Bozeman, MT, USA). To synchronize electrophysiological and behavioral recordings, a 0.5 Hz TTL pulse was sent from an Arduino to both the Neuropixels and Digital Lynx systems. One female rat was excluded from analysis due to probe failure. Therefore, data from 9 out of 10 implanted animals were included in the analyses of recording data.

### Histology, probe tracking and cell sorting

Following the completion of all recordings, animals were first anesthetized with 2-3% isoflurane (in oxygen at 1 L/min) to allow for extraction of the Neuropixels probes. They were then euthanized with sodium pentobarbital. Transcardial perfusion was performed using phosphate-buffered saline (1 × PBS, 0.1 M, pH 7.4), followed by 4% paraformaldehyde in 0.1 M PBS. Brains were post-fixed in 4% paraformaldehyde and subsequently cryoprotected in a 30% sucrose solution. Coronal brain sections (40 µm thick) were prepared and mounted on glass slides.

Prior to implantation, Neuropixels probes were coated with 2% DiI (D282, Thermo Fisher) dissolved in pure isopropyl alcohol to enable post hoc tracing of the probe tracks. After sectioning, DAPI was applied to the mounted slices before cover slipping. Fluorescent images were acquired using an Olympus slide scanner (VS200, Olympus) at emission wavelengths of 455 nm for DAPI and 565 nm for DiI. Recording sites with clear theta-band activity and located within 2 mm of the dorsal border of mEC were classified as dorsal mEC; more ventral sites were categorized as ventral mEC.

Spike sorting was carried out using Kilosort 2.0 (MouseLand/Kilosort) with the following parameters: ops.minfr_goodchannels = 0.01, ops.Th = [10 4], and ops.minFR = 1/1000. Manual curation of clusters was performed in Phy (cortex-lab/phy). Units were classified as PNs if their average spike waveform width was ≥ 0.4 ms.

### Local field potential analysis

To analyze theta oscillations during each recording session, one LFP channel was selected from the sites confirmed to be located in mEC based on the histology and theta amplitude. To detect theta episodes, theta power was computed across whole recording sessions, including open box and sleep sessions and 80 trials of the alternation task. Theta episodes were defined as periods when z-scored theta power was >0 for at least 0.5 s. Two theta episodes were joined if their gap was ≤0.5 s. The initial theta bout length in the delay was determined by selecting the first theta episode end time in the delay zone, or if there was no theta detected, when the animal entered the delay zone by setting the initial theta bout length to 0 s.

### Low-dimensional visualization of population activity

To visualize the low-dimensional structure of mEC population activity in the WM task, spike trains of PNs were discretized into 200-ms time bins. Binned spike trains of each neuron were smoothed using a Gaussian kernel with 400-ms width and were then z-scored across time to normalize firing-rate variability. For each bin, the animal’s position on the maze was assigned based on the corresponding behavioral data. The resulting population activity matrix (time bins × neurons) was embedded into a low-dimensional space using Uniform Manifold Approximation and Projection (UMAP) ^69^. Each point in the low-dimensional embedding represents the population firing pattern within a 200-ms time bin during the session. UMAP was performed with the following parameters: n_neighbors = 20, min_dist = 0.2, spread = 2.0, metric = ‘cosine’, method = ‘mex’, and n_components = 3.

### Definition of time cells and distance cells

Spike trains obtained from individual neurons were aligned by using the time when the animal was within 15 cm of the delay barrier (shown in **Figure 1**) as the delay onset. The temporal firing profile in the delay area was calculated by binning spikes into 150-ms windows followed by a Gaussian kernel convolution with a sigma of 300 ms.

For each neuron that had spikes in the delay area in more than 10 trials of any of the four conditions (from 20 trials per condition, obtained by combining the two 10-trial blocks in identical conditions), temporal stability was assessed by first calculating the Pearson correlation between every pair of trials with spikes (10-20 trials resulted in 45-190 correlation values) and by then taking the median of all correlation values. A shuffled stability distribution was calculated for each neuron by randomly shifting spike trains in the delay interval for one of the trials in each pair (by at least 500 ms and with circularly wrapping the values). As for the actual data, the median of all pairwise comparisons was then taken. The shuffling procedure was repeated 1000 times, and the cell was considered a time cell in a delay/treadmill condition if the actual stability exceeded the 95th percentile of the shuffled distribution.

In sessions where treadmill speed was varied during the delay period, spike trains were also aligned to the distance traveled on the treadmill belt, starting from the moment the animal reached the delay barrier. Distance-based firing profiles were computed by binning spikes into 2 cm intervals and smoothing with a Gaussian kernel (σ = 4 cm). As with temporal analysis, neurons with spiking activity in more than 10 trials for any of the four conditions were analyzed for distance stability. Pearson correlations were calculated for all pairs of valid trials, and the median correlation value was taken as the stability metric. The shuffling procedure was performed as for time cells, but with circular shifts by at least 500 ms × treadmill speed. A neuron was classified as a distance cell in the treadmill-on delay condition if its actual stability exceeded the 95^th^ percentile of the corresponding shuffled distribution. If a neuron met criteria for both time cell and distance cell classification, it was assigned to the category (time or distance) corresponding to the higher median trial-to-trial correlation value.

### Grid cell identification

Rats foraged freely for randomly scattered drops of chocolate milk in a square open-field environment (1.2 m × 1.2 m) with black walls and a white cue card fixed to one wall. Each session consisted of three 10-minute foraging blocks, separated by 5-min rest periods in a familiar sleep box. Spatial firing rate maps were constructed for each 10-min block and for the combined data across all three blocks. Position data were binned into 3 cm × 3 cm spatial bins, and firing rates were smoothed using a Gaussian kernel with a 3 cm standard deviation. Two-dimensional spatial autocorrelograms were computed from the smoothed rate maps to quantify periodicity in firing patterns.

Gridness scores were calculated from the spatial autocorrelograms by measuring the degree of hexagonal symmetry. Specifically, the autocorrelogram was rotated by 30°, 60°, 90°, 120°, and 150°, and the Pearson correlation between the original and rotated maps was computed. The gridness score was defined as the minimum difference between the average correlations at 60° and 120° (expected to be high for hexagonal symmetry) and the average of the correlations at 30°, 90°, and 150° (expected to be low). A PN was classified as a grid cell if the gridness score exceeded 0.3 in any individual 10-minute block or in the combined map across all three blocks.

### Definition of periodic cells

To evaluate the distance periodicity of single neurons during treadmill-on delay intervals, firing rates were computed as a function of distance traveled, with spikes binned in 3-cm intervals starting from the moment the animal reached the delay barrier, with a gaussian smoothing kernel of 6 cm. An autocorrelation of the distance-binned firing rate was then computed for each neuron, using lags up to the maximum distance traveled during the delay. To assess whether a PN’s firing exhibited significant distance periodicity, a null distribution was generated through a blockwise shuffling procedure. Specifically, spike bins within the delay interval were randomly shifted, preserving firing statistics while disrupting long-range structure. For the shuffled instances, autocorrelograms were computed, and the 99% confidence threshold was defined as the mean plus 2.58 standard deviations of the shuffled autocorrelogram. Peaks in the empirical autocorrelogram were identified using MATLAB’s findpeaks function. A neuron was classified as a distance-periodic cell if it exhibited at least one peak that met the following criteria: (1) peak amplitude ≥ 0.15, (2) peak-to-trough amplitude ≥ 0.1, and (3) peak height exceeded the 99% confidence threshold derived from the shuffled data. The lag corresponding to the first significant peak was taken as the PN’s distance periodicity.

### Identification of delay sequences

To identify temporally structured neuronal activity patterns during delay periods, we extracted neuronal sequences from mEC PN rate matrices using sequential non-negative matrix factorization (seqNMF), adapted from ref. 70 and implemented via the publicly available seqNMF toolbox (https://github.com/FeeLab/seqNMF).

#### Spike preprocessing

Spike trains from putative PNs were first discretized into a time-by-neuron matrix using 200-ms non-overlapping bins. The binned firing rates were then smoothed using a Gaussian kernel with a sigma of 400 ms to reduce variability. Following smoothing, each PN’s activity was z-scored across the session. To reduce the influence of low firing-rate noise, bins with z-scores below zero were set to zero.

#### Selection of behavioral epochs

Analysis was restricted to data recorded during treadmill-on blocks, only when the animal was in the delay intervals. We intentionally excluded the return arms, stem and T zone—despite their continuous running structure—to avoid identifying sequences that primarily reflect repeated spatial trajectories rather than temporal organization during the delay.

#### Sequence detection with seqNMF

SeqNMF was applied to the z-scored, zero-corrected rate matrix extracted from the delay epochs. The algorithm identifies repeated, temporally structured patterns of co-activation across neurons. Formally, given a data matrix *X* of size *n* × *t* (with *n* neurons and *t* time bins), seqNMF factorizes *X* as the matrix product:

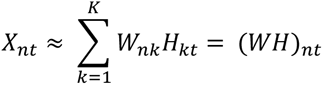

where *X_nt_* is the activity of neuron *n* at time *t*, *W_nk_* represents the contribution of neuron *n* to the *k*-th sequence pattern, and *H_kt_* is the activation strength of the *k*-th sequence at time *t*. In this formulation, *W* reflects the contribution of neuron *n* to the *k*-th sequence pattern, while *H* captures the temporal dynamics of when each sequence is expressed. We used parameters chosen to capture the behavioral timescale of delay activity: a time bin size of 200 ms, number of factors *k* = 8, regularization strength *λ* = 0.001, and sequence length *ℓ* = 15 or *ℓ* = 25, corresponding to durations of 3.0 s or 5.0 s, respectively. The choice of *ℓ* was guided by the average temporal spacing of periodic cell firing peaks within each recording session, ensuring that the extracted sequences approximately matched the underlying periodicity of the mEC cell population.

#### Cell inclusion criteria for sequence components

To determine whether a neuron significantly contributed to a detected sequence in the weight matrix *W_nk_*, we established a statistical threshold using a surrogate data approach designed to preserve each cell’s intrinsic temporal firing structure during the delay intervals. Specifically, following ref. 71 for each PN, we generated a set of synthetic spike trains that preserved both the neuron’s spike count and its spike-time autocorrelogram. This ensured that key features of the neuron’s intrinsic firing dynamics were maintained, while eliminating structured co-activation across the neuron population.

The same seqNMF analysis as used for the original data was applied to each of 100 independently generated surrogate datasets. This yielded a null distribution of weights for each neuron across all sequence components, resulting in 800 shuffled *W* values per neuron (8 sequences × 100 shuffle instances). A neuron was considered to be contributing significantly to a sequence if its actual weight in a given sequence exceeded its own 99^th^ percentile threshold from shuffled data. Additionally, any time bin in which more than five neurons exhibited peak activation led to the exclusion of all neurons peaking in that bin.

Only sequences that included ≥10 significantly contributing neurons were retained for subsequent analyses. After identifying the engaged cells, the corresponding temporal activation pattern *H_kt_* was recalculated using the pseudo-inverse as: *H* ≈ *X*/*W*. To reduce noise, a low-pass Butterworth filter was then applied to *H*. Given the 200-ms time bin resolution, the sampling frequency was set to *f_s_* = 5 Hz and the cutoff frequency to *f_c_* = 0.5 Hz. The normalized cutoff frequency was computed as *W_n_* = *f_c_* / (*f_s_*/2). A 4^th^-order low-pass Butterworth filter was designed using these parameters with the MATLAB command [b, a] = butter(4, *W_n_*, ’low’).

Candidate sequence events were identified by detecting peaks in the filtered *H* signal using MATLAB’s findpeaks function. For each detected peak, the event window was defined as the interval between the nearest troughs before and after the peak. To assess the statistical significance of these candidate events, a two-step shuffle procedure was performed. The original data matrix *X* was first shuffled by randomly permutating the order of cells, and then by shuffling the time bins. The sequence amplitude *H* calculated from the shuffled matrix was then subjected to the same filtering and peak-detection procedure as *H* calculated from the original data. The 95^th^ percentile of the peak values from the shuffled data was used as the threshold for the event detection. Candidate events with peak amplitudes in the original *H* exceeding this threshold were classified as significant, and their corresponding trough-to-trough intervals were retained as sequence events. The peak amplitude time of significant events was used to calculate autocorrelograms for each sequence and crosscorrelograms across different sequences. To assess how sequence participation scaled with the size of the recorded population, we calculated the fraction of mEC PNs that participated in at least one sequence, relative to the total number of recorded PNs. This relationship was modeled using a nonlinear saturation fit, and the plateau value was reported for statistically significant fits (p < 0.05).

#### Sequence order correlation with neural activity in other maze segments

To examine whether the order of neurons within treadmill-on delay sequences (detected with seqNMF) reflected spatial firing patterns elsewhere on the maze, we compared the order of neurons in seqNMF-detected sequences to the spatial order of the same neurons during running in other maze segments. Spatial order was assessed within fixed-length trajectory windows of 50 cm, 80 cm, and 110 cm, advanced in 10-cm increments along each trajectory. For each neuron that was part of a seqNMF-detected sequence, we calculated the mean spike location in the windows along the return arm and the stem-to-reward zone in the forward and reverse order and ranked the neurons accordingly. The rank order in trajectory windows was then compared to the order in seqNMF-detected sequences (as defined by the component’s weight vector *W_nk_*) using Spearman’s rank correlation and considered significantly correlated if p < 0.05.

During treadmill-off delay intervals, population events were identified as epochs when the population firing rate of PNs that were included in a seqNMF-detected sequence exceeded the mean population firing rate by four standard deviations in consecutive 10-ms bins. The order of the neurons’ average spike times within each population event was then compared to the order in the seqNMF-detected sequence using Spearman’s rank correlation and considered significantly correlated if p < 0.05.

### Trajectory-selective cells and sequences

To identify trajectory-selective activity patterns of individual cells, we computed a normalized firing rate difference for each PN during each delay condition in 5-s time intervals:

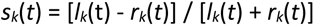

where *s_k_*(*t*) is the normalized rate difference between upcoming left-turn or right-turn trials for neuron *k* at the time interval *t*, and *ℓ* and *r* are the mean firing rates during upcoming left-turn and right-turn trials. Neurons with mean firing rates below 0.5 Hz in the region of interest were excluded. Significance was assessed by generating a null distribution from 1000 random permutations of trial labels. A neuron was considered trajectory-selective in a 5-s interval during the delay if its observed *s_k_*(*t*) exceeded the 95^th^ percentile of the shuffled distribution. To quantify the proportion of trajectory-selective neurons, we included only 20-trial blocks (e.g., treadmill-on/30-s) with ≤5 error trials. The low number of errors implies that the return arm and upcoming turn were opposite (e.g., left return and upcoming right turn) for most trials.

To distinguish return-selective from upcoming turn-selective cells, we analyzed 20-trial blocks with ≥6 error trials. Selectivity indices for return-selective cells were computed using left- vs. right-return trials preceding the delay, and indices for upcoming turn-selective cells were computed using left- vs. right-turn trials following the delay. Neurons were classified as return-selective or upcoming turn-selective if the selectivity index exceeded the 95^th^ percentile of the corresponding shuffled distribution. To avoid category overlap, any neuron that met criteria for both return- and turn-selectivity was excluded from both classifications. The proportions of return-selective neurons and the proportions of upcoming turn-selective neurons were then compared to chance levels.

Similar analyses were performed to determine whether sequences during treadmill-on delay intervals show memory-related coding. For calculations of differences, the sequence amplitude *H* in every 10-s interval was used, and the analysis was performed for session blocks with different error thresholds (≥3, 4, 5 and 6). Sequences were classified as return-selective or upcoming turn-selective if their selectivity index exceeded the 95^th^ percentile of the corresponding shuffled distribution. In addition, sequence amplitudes were also compared between trials with correct or incorrect upcoming choices (following the delay) and classified as outcome-selective if the actual amplitude difference exceeded 95^th^ percentile of the corresponding shuffled distribution.

### Statistics

One-way ANOVA was used to analyze the behavioral performance (**Figure 1a**), theta power and length (**Figure 1b**, **Figure S1d**), and sequence event counts and amplitude (**Figure 7c**). Two-way ANOVA was used to analyze the time/distance cells under different delay conditions (**Figure 3c**) and to compare the proportions of distance and time cells (**Figure S5e**). T-tests were used to compare the proportion of periodic cells (**Figure 5c**), the theta power across the figure-8 maze (**Figure S1g**) and the spatial stability (**Figure S2c**). Spearman’s correlations were used to analyze the relationship between grid distance and periodic distance (**Figure 5e**), the correlation of average rate in different maze sections (**Figure S4**), the correspondence of sequence order across maze segments (**Figure 6e** and **f**, **Figures S8-S10**, **Figure S13**) and the relationships between cell numbers and sequence properties (**Figure S11a-c**). Non-linear saturation fitting was used to check whether a plateau existed for the sequence cell participation (**Figure 6b**). Binomial tests were used to compare turn-selective single cell and sequence proportions to the 5% chance level (**Figure 4b** and **c**, **Figure 8c**, **Figure S14**). Average values correspond to the mean ± SEM unless indicated otherwise. All statistical significance was set to a level of 0.05, divided among two sides, except that binomial tests were one-sided. Post-hoc adjustments for multiple comparisons were performed.

