## Supplemental Material for "Working memory retention by medial entorhinal cortex low-dimensional neural dynamics"

### Figure S1

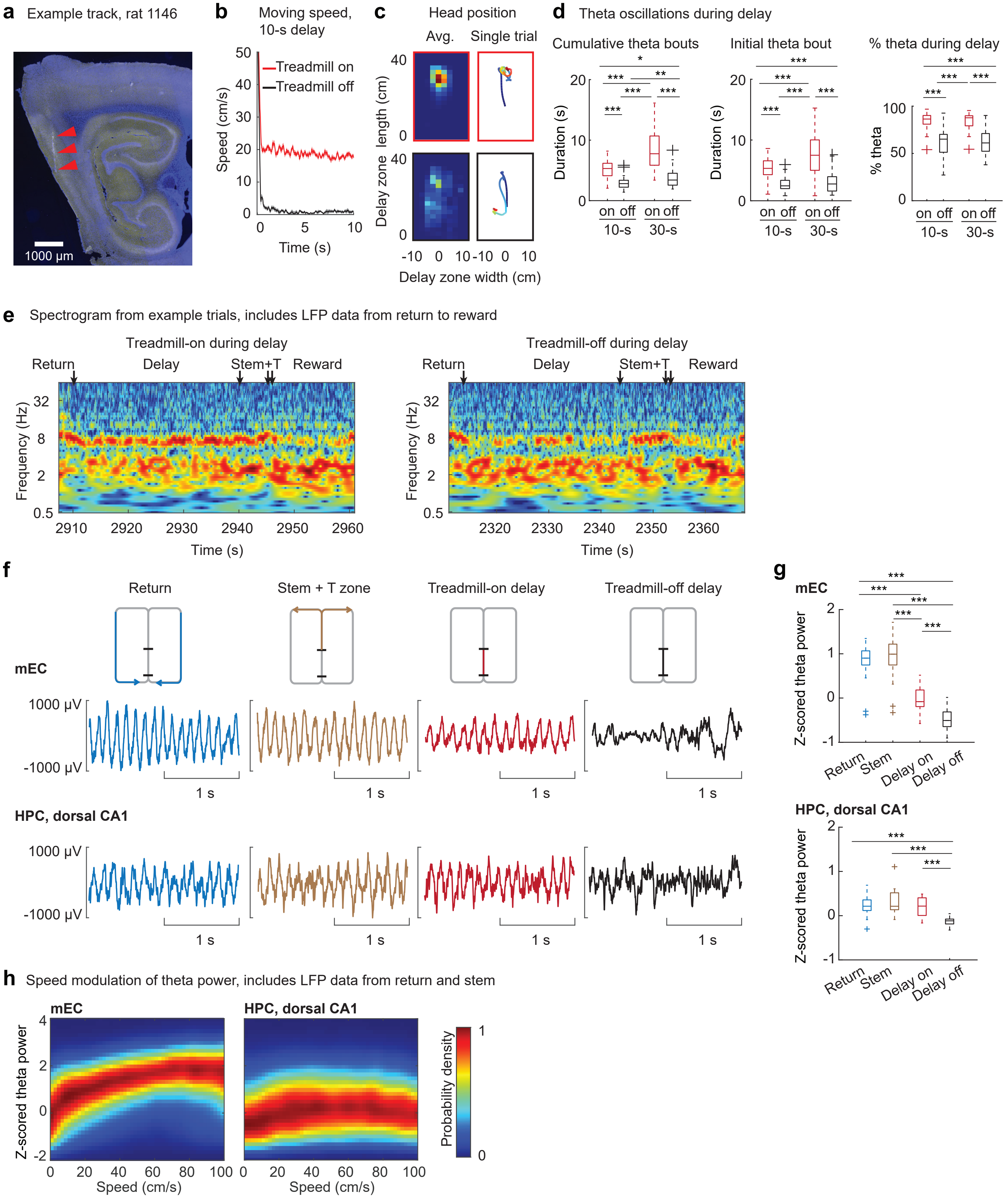

a Example track, rat 1146
1000 µm
b Moving speed, 10-s delay
Treadmill on
Treadmill off
Speed (cm/s)
Time (s)
c Head position
Avg.
Single trial
Delay zone length (cm)
Delay zone width (cm)
d Theta oscillations during delay
Cumulative theta bouts
Initial theta bout
% theta during delay
Duration (s)
% theta
on off
10-s
30-s
e Spectrogram from example trials, includes LFP data from return to reward
Treadmill-on during delay
Treadmill-off during delay
Return
Delay
Stem+T
Reward
Frequency (Hz)
Time (s)
f
Return
Stem + T zone
Treadmill-on delay
Treadmill-off delay
mEC
1000 µV
-1000 µV
1 s
HPC, dorsal CA1
g
mEC
Z-scored theta power
Return
Stem
Delay on
Delay off
HPC, dorsal CA1
h Speed modulation of theta power, includes LFP data from return and stem
mEC
HPC, dorsal CA1
Z-scored theta power
Speed (cm/s)
Probability density

#### Figure S1, continued

**Figure S1. Histology, behavior and theta oscillations.** **a**, Track from one of the four shanks of a Neuropixels 2.0 probe, shown in a sagittal section of an example rat (rat 1146). The probe track traverses medial entorhinal cortex (mEC) superficial layers. Red arrows point to the track through mEC. **b**, Mean running speed during 10-s treadmill-on (red) and treadmill-off (black) delay intervals from one example rat (rat 1138, averaged over all trials from 4 recording days). **c**, Head position during delay intervals, from the same rat as in a. Top, treadmill-on trials (red frame); bottom, treadmill-off trials (black frame). Left, average occupancy across trials (red, maximum occupancy time; blue, unvisited locations). Right, example single-trial trajectory, color scale depicts start (blue) to end (red) of delay. **d**, Left: Average theta bout duration was longer in treadmill-on compared to off conditions (n = 33 sessions; $F(3,128) = 53.72$, $p = 1.5 \times 10^{-22}$, one-way ANOVA; Tukey–Kramer post hoc tests: on/10-s vs. off/10-s: $p = 4.2 \times 10^{-5}$; on/10-s vs. on/30-s: $p = 1.1 \times 10^{-8}$; off/10-s vs. off/30-s: $p = 0.475$; on/30-s vs. off/30-s: $p = 3.8 \times 10^{-9}$). Center: The duration of the initial theta bout during the delay was longer in treadmill-on compared to off conditions (n = 33 sessions; $F(3,128) = 33.35$, $p = 5.4 \times 10^{-16}$, one-way ANOVA; Tukey–Kramer post hoc tests: on/10-s vs. off/10-s: $p = 6.9 \times 10^{-5}$, on/10-s vs. on/30-s: $p = 6.5 \times 10^{-5}$, off/10-s vs. off/30-s: $p = 0.97$, on/30-s vs. off/30-s: $p = 3.8 \times 10^{-9}$). Right, Percentage of time in delay period with high theta power was higher in treadmill-on compared to off conditions (n = 33 sessions; $F(3,128) = 46.63$, $p = 2 \times 10^{-20}$, one-way ANOVA; Tukey–Kramer post hoc tests: on/10-s vs. off/10-s: $p = 3.8 \times 10^{-9}$, on/10-s vs. on/30-s: $p = 1$, off/10-s vs. off/30-s: $p = 0.99$, on/30-s vs. off/30-s: $p = 3.8 \times 10^{-9}$). **e**, Time-frequency spectrograms of mEC LFP from example trials with the treadmill on and with the treadmill off during 10-s delay intervals. Maze segments are indicated on top. **f**, Example raw LFP traces from mEC (top) and dorsal hippocampal CA1 (bottom) during return arm, stem + T zone, treadmill-on delay and treadmill-off delay. For comparison to mEC, hippocampal data from ref. 26 were reanalyzed. **g**, Top, mEC theta power was higher in the return arm and stem compared to the treadmill-on delay and higher in the treadmill-on delay than in treadmill-off delay (n = 33 sessions; return vs. treadmill-on: $p = 2 \times 10^{-9}$, return vs. treadmill-off: $p = 1 \times 10^{-14}$, stem vs. treadmill-on: $p = 5 \times 10^{-10}$, stem vs. treadmill-off: $p = 5 \times 10^{-14}$, treadmill-on vs. treadmill-off: $p = 3 \times 10^{-9}$; all other comparisons, $p > 0.05$, Bonferroni corrected paired t-tests). The lower theta power at treadmill speeds (10-20 cm/s) is consistent with the finding that mEC peak power is not reached until speeds exceed 60 cm/s. Bottom: Theta power in hippocampus (HPC) was lower in treadmill-off than in all other maze zones, while the other maze zones did not differ from each other (n = 16 sessions; return vs. treadmill-off: $p = 7 \times 10^{-5}$, stem vs. treadmill-off: $p = 1 \times 10^{-5}$, treadmill-on vs. treadmill-off: $p = 0.0001$, all other comparisons: $p > 0.05$, Bonferroni corrected paired t-tests). **h**, Probability density of theta power measured in mEC (left) and in the hippocampal CA1 area (right) during return and stem segments. Theta power within each brain region was z-scored across all trials. Running speed was divided into 2 cm/s bins, and zero to maximum probability of z-scored theta power within each speed bin is depicted as in the color bar to the right. Box plots: central line, edges, whiskers and plus signs; median, the 25th/75th percentile, maximum/minimum and outliers. * $p < 0.05$, ** $p < 0.01$, *** $p < 0.001$.

#### Figure S2

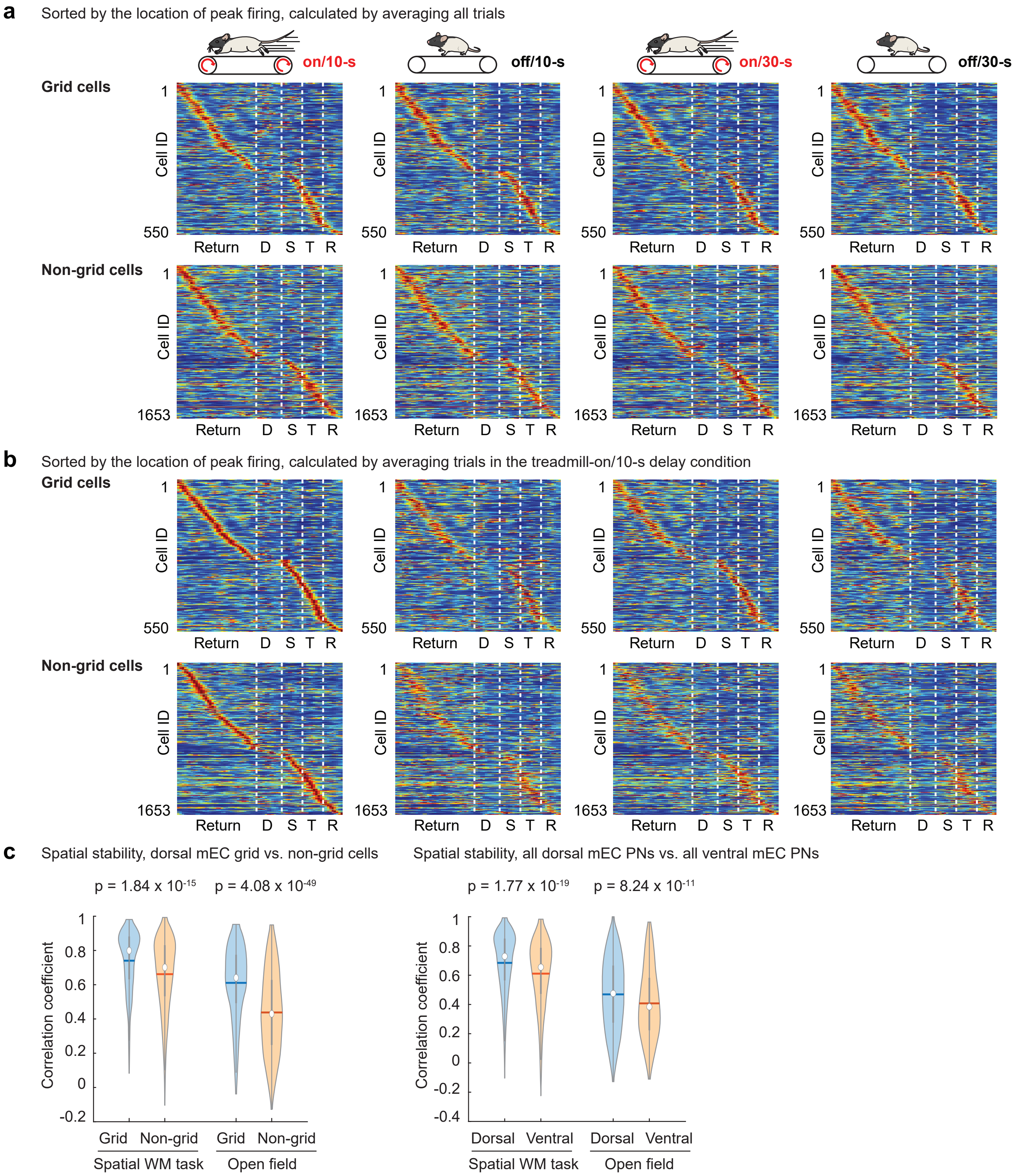


**Figure S2. Spatial firing patterns and spatial stability of mEC cells in the spatial WM task. a**, Spatial firing patterns of dorsal mEC grid cells (top) and non-grid cells (bottom) in all maze zones. Cells are sorted by the location of peak firing, calculated from the average firing rate across all four treadmill/delay conditions. To ensure robust classification of grid cells, only recording sessions with good coverage in the open-field are included. **b**, Same data as in a, with cells sorted by the location of peak firing during treadmill on/10-s delay trials. **c**, Left: Spatial stability of grid cells was higher than stability of non-grid cells (spatial WM task: $p = 2 \times 10^{-15}$; open field: $p = 4 \times 10^{-49}$, n = 550 grid and 1653 non-grid cells, two-sided t test). Right: Spatial stability of dorsal mEC PNs was higher than spatial stability of ventral mEC PNs (spatial WM task: $p = 2 \times 10^{-19}$; open field: $p = 8 \times 10^{-11}$, n = 2642 dorsal and 891 ventral cells, two-sided t-test). D, Delay; S, Stem; T, T zone; R, Reward.

### Figure S3

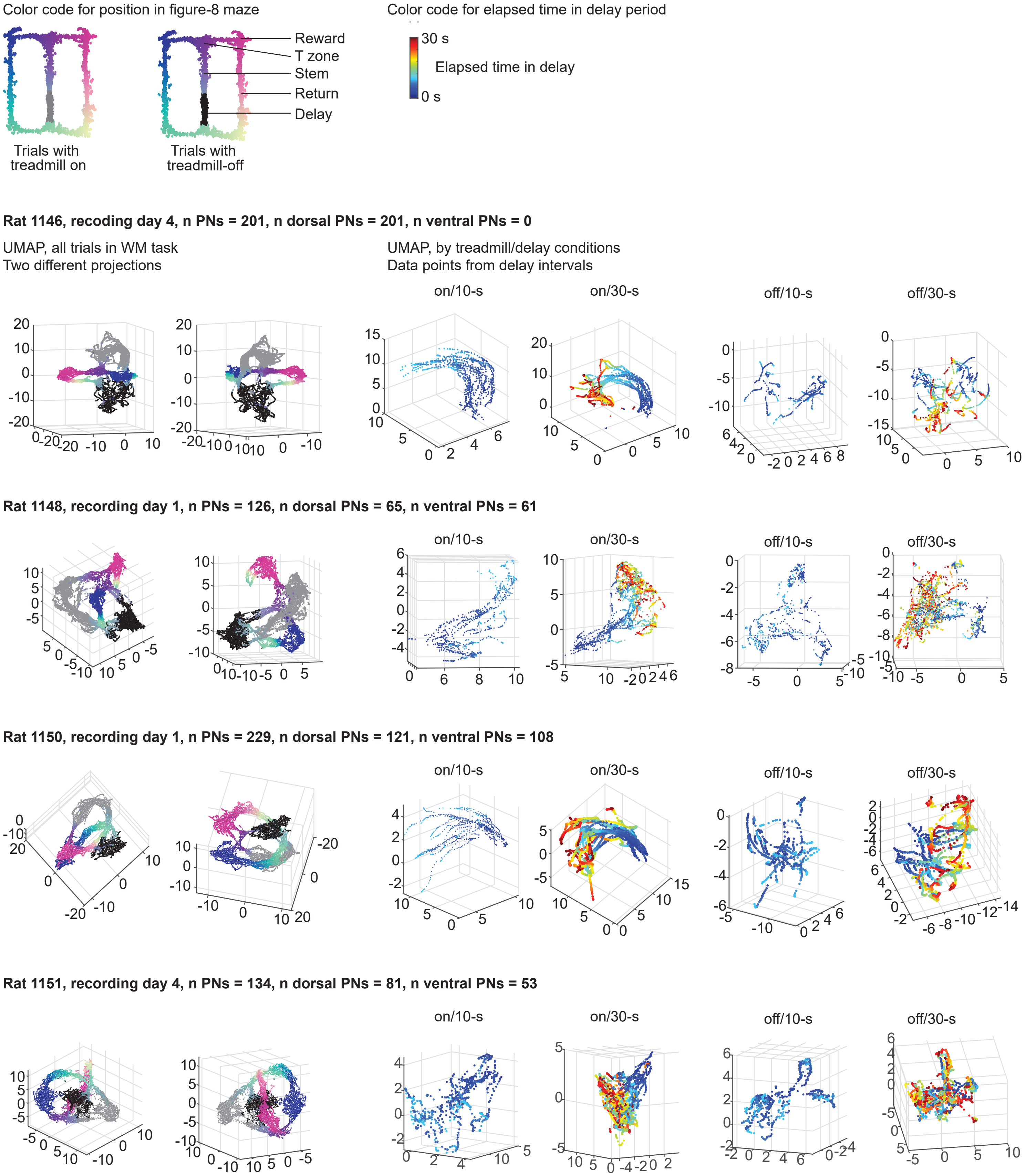

Color code for position in figure-8 maze
Reward
T zone
Stem
Return
Delay
Trials with treadmill on
Trials with treadmill-off
Color code for elapsed time in delay period
30 s
Elapsed time in delay
0 s
Rat 1146, recoding day 4, n PNs = 201, n dorsal PNs = 201, n ventral PNs = 0
UMAP, all trials in WM task
Two different projections
UMAP, by treadmill/delay conditions
Data points from delay intervals
on/10-s
on/30-s
off/10-s
off/30-s
Rat 1148, recording day 1, n PNs = 126, n dorsal PNs = 65, n ventral PNs = 61
on/10-s
on/30-s
off/10-s
off/30-s
Rat 1150, recording day 1, n PNs = 229, n dorsal PNs = 121, n ventral PNs = 108
on/10-s
on/30-s
off/10-s
off/30-s
Rat 1151, recording day 4, n PNs = 134, n dorsal PNs = 81, n ventral PNs = 53
on/10-s
on/30-s
off/10-s
off/30-s

#### Figure S3, continued

**Rat 1137, recording day 1, n PNs = 95, n dorsal PNs = 4, n ventral PNs = 91**

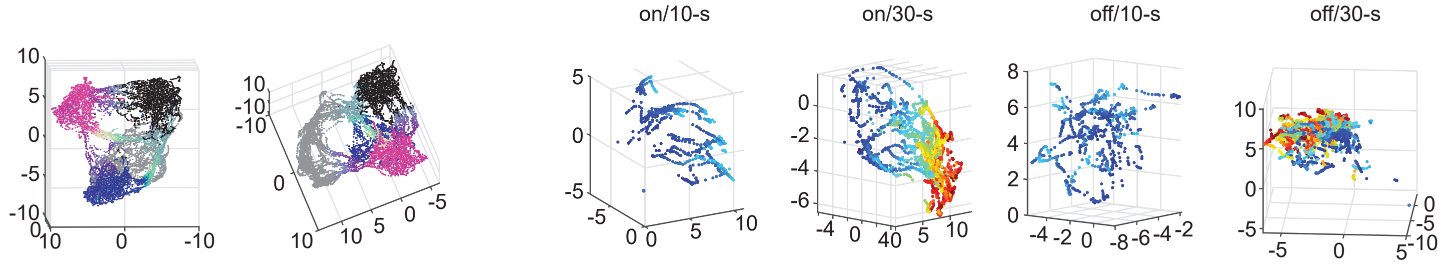


**Rat 1140, day 1 recording, n PNs = 121, n dorsal PNs = 38, n ventral PNs = 83**

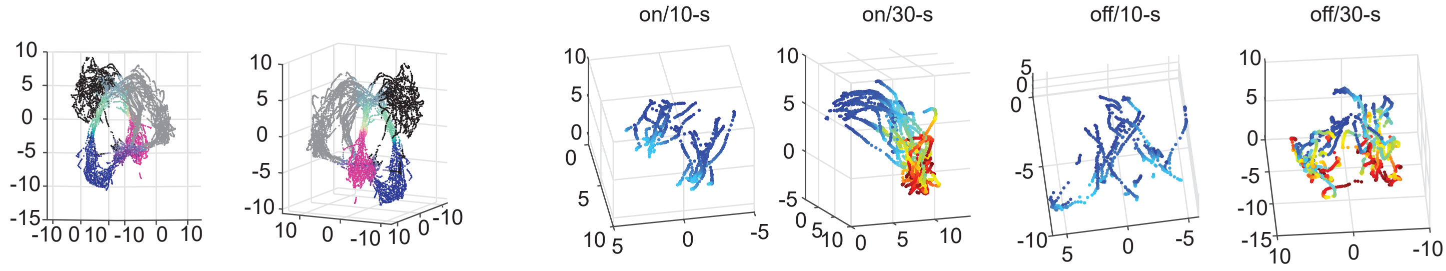


**Rat 1138, recording day 2, n PNs = 171, n dorsal PNs = 171, n ventral PNs = 0**

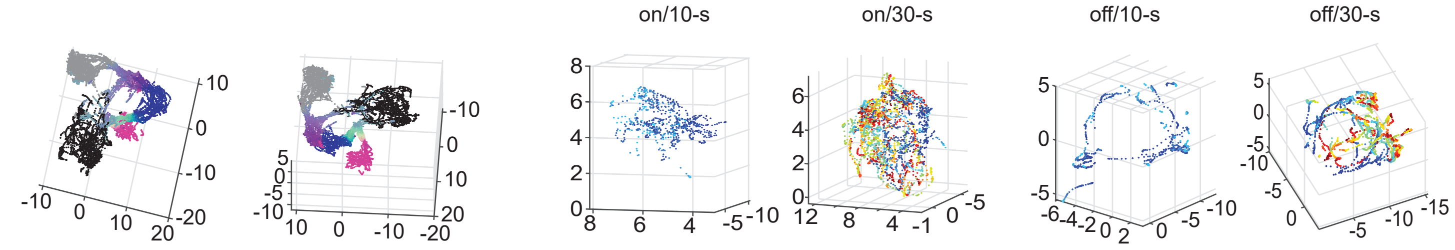


**Rat 1139, recording day 2, n PNs = 75, n dorsal PNs = 75, n ventral PNs = 0**

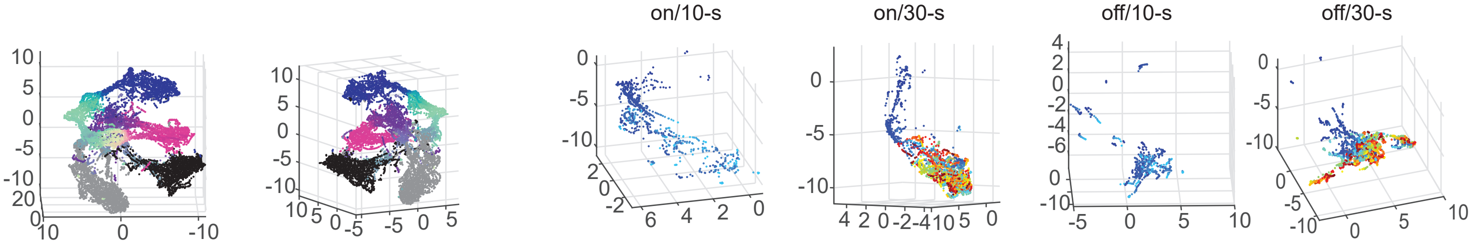


**Rat 1147, recording day 3, n PNs = 57, n dorsal PNs = 57, n ventral PNs = 0**

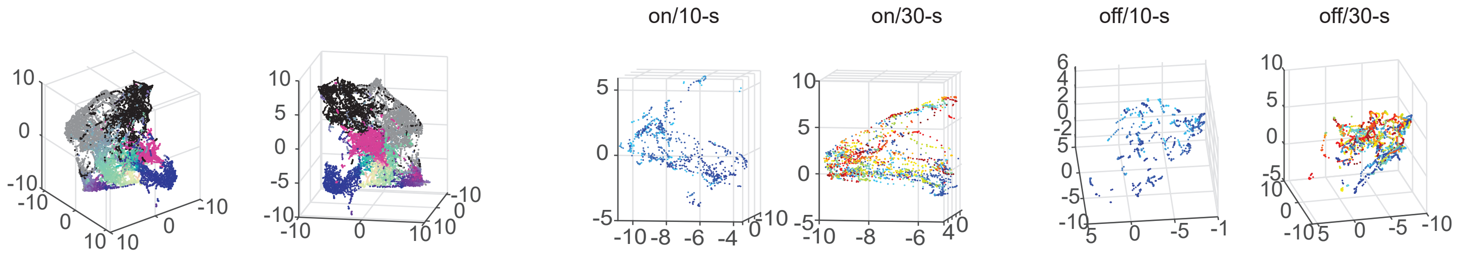


**Figure S3. Low dimensional embeddings of mEC population activity resembled maze structure and differed across delay conditions.** UMAP projections of simultaneously recorded mEC cells in the WM task are shown for one recording session from each rat. Spatial positions in the figure-8 maze and elapsed time in the delay interval are color coded according to the schematics on top. In projections of the entire figure-8 maze, gray is used for the delay interval in treadmill-on trials and black is used for the delay in treadmill-off trials. For each rat, the two leftmost panels show the same embedding viewed from different angles. The four panels on the right display neural activity trajectories from trials with treadmill-on/10-s delays, treadmill-on/30-s delays, treadmill-off/10-s delays and treadmill-off/30-s delays, respectively, projected into the same low-dimensional space as the left panels.

### Figure S4

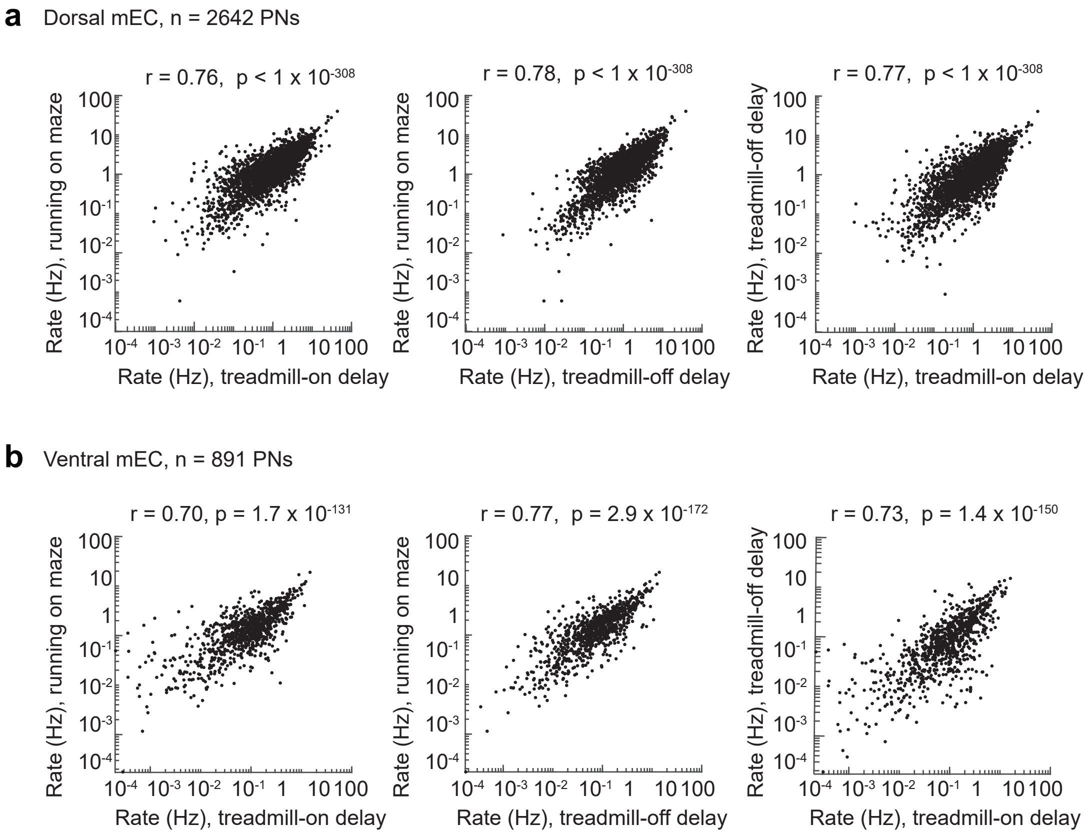


**Figure S4. Average firing rates during treadmill-on and treadmill-off delay intervals were strongly correlated with each other and with average firing rates during maze running.** **a**, Dorsal mEC neurons. Firing rates during each of the delay conditions and during running on the maze were correlated (treadmill-on vs. maze: Spearman's r = 0.76, $p < 1 \times 10^{-308}$; treadmill-off vs. maze: r = 0.78, $p < 1 \times 10^{-308}$; treadmill-on vs. treadmill-off: r = 0.77, $p < 1 \times 10^{-308}$, n = 2642 PNs). **b**, Ventral mEC neurons. Firing rates during each of the delay conditions and during running on the maze were correlated (treadmill-on vs. maze: r = 0.70, $p = 1.7 \times 10^{-131}$; treadmill-off vs. maze: r = 0.77, $p = 2.9 \times 10^{-172}$; treadmill-on vs. treadmill-off: r = 0.73, $p = 1.4 \times 10^{-150}$, n = 891 PNs).

#### Figure S5

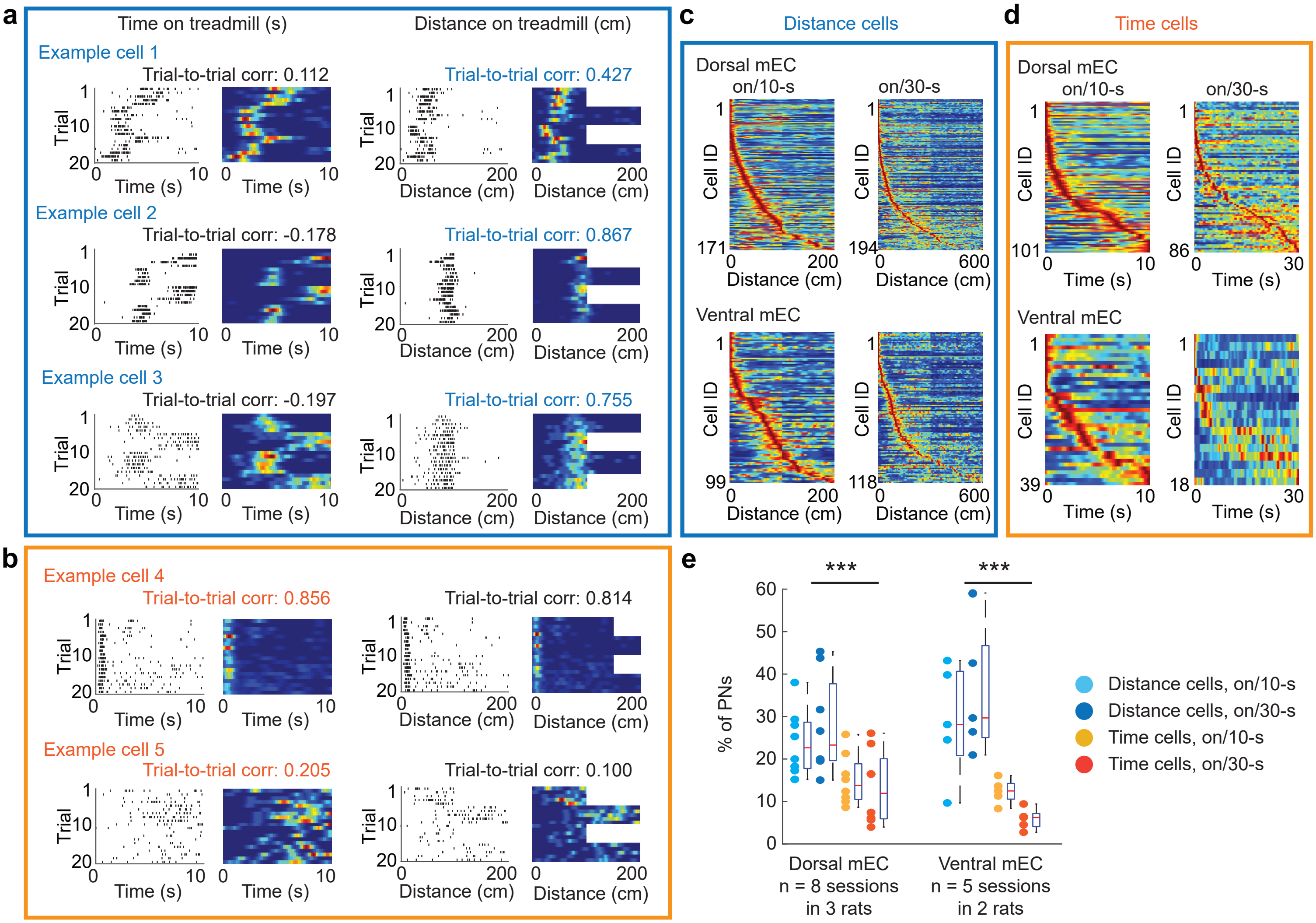


**Figure S5. In mEC cells, distance coding was more pronounced than time coding during treadmill-on delay intervals. a**, Example cells that coded for distance rather than time. Each row shows a single mEC neuron. Firing patterns for each trial are shown by plotting each spike (black ticks) or by color coding the firing rate (red, maximum; blue, 0 Hz). Median trial-to-trial correlations are indicated on top of each set of panels. **b**, Example cells that coded for time rather than distance, plotted as in a. **c**, Average firing rate of each distance cell (one per row) in dorsal (top) and ventral (bottom) mEC during 10-s (left) and 30-s (right) delay intervals. Cells are ordered by the distance of the firing peak within each condition. **d**, Average firing rate of each time cell (one per row) in dorsal (top) and ventral (bottom) mEC during 10-s (left) and 30-s (right) delay intervals. Cells are ordered by the time of the firing peak within each condition. **e**, A larger proportion of mEC cells coded for distance compared to time (dorsal mEC: n = 8 sessions from 3 animals, 10-s vs. 30-s: $F(1,28) = 0.099$, $p = 0.756$, distance vs. time: $F(1,28) = 14.35$, $p = 0.0007$; interaction: $F(1,28) = 0.84$, $p = 0.37$; two-way ANOVA; ventral mEC: n = 5 sessions from 2 animals; 10-s vs. 30-s: $F(1,16) = 0.0004$, $p = 0.984$; distance vs. time: $F(1,16) = 25.42$, $p = 0.0001$, interaction: $F(1,16) = 2.04$, $p = 0.17$; two-way ANOVA). *** $p < 0.001$.

#### Figure S6

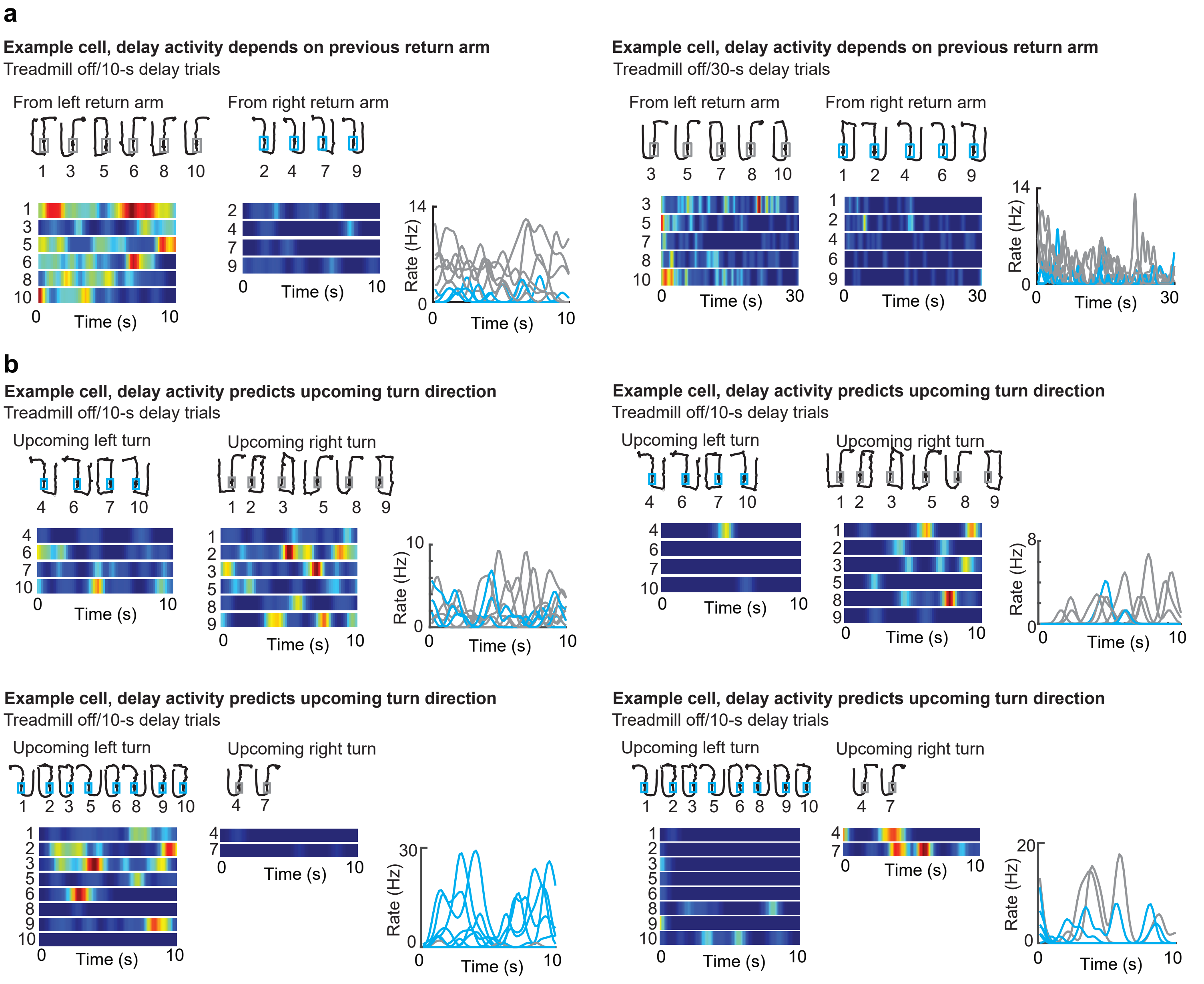


**Figure S6. Delay-period activity in a subset of mEC cells was memory-related, in particular with the treadmill off. a**, Two example mEC neurons with higher firing rates during the delay intervals after returning from the left-arm. Top of panel, rat trajectories for each trial (black) with the delay zone highlighted (gray box, from left return arm; blue box, from right return arm). Bottom of panel, firing rate throughout the delay is color coded (red, maximum; blue, 0 Hz). Right of panel, trial-wise firing rate throughout delay intervals (gray lines, from left return arm; blue lines, from right return). **b**, Four example mEC neurons with higher delay-period activity for one of the upcoming turn directions, depicted as in a.

#### Figure S7

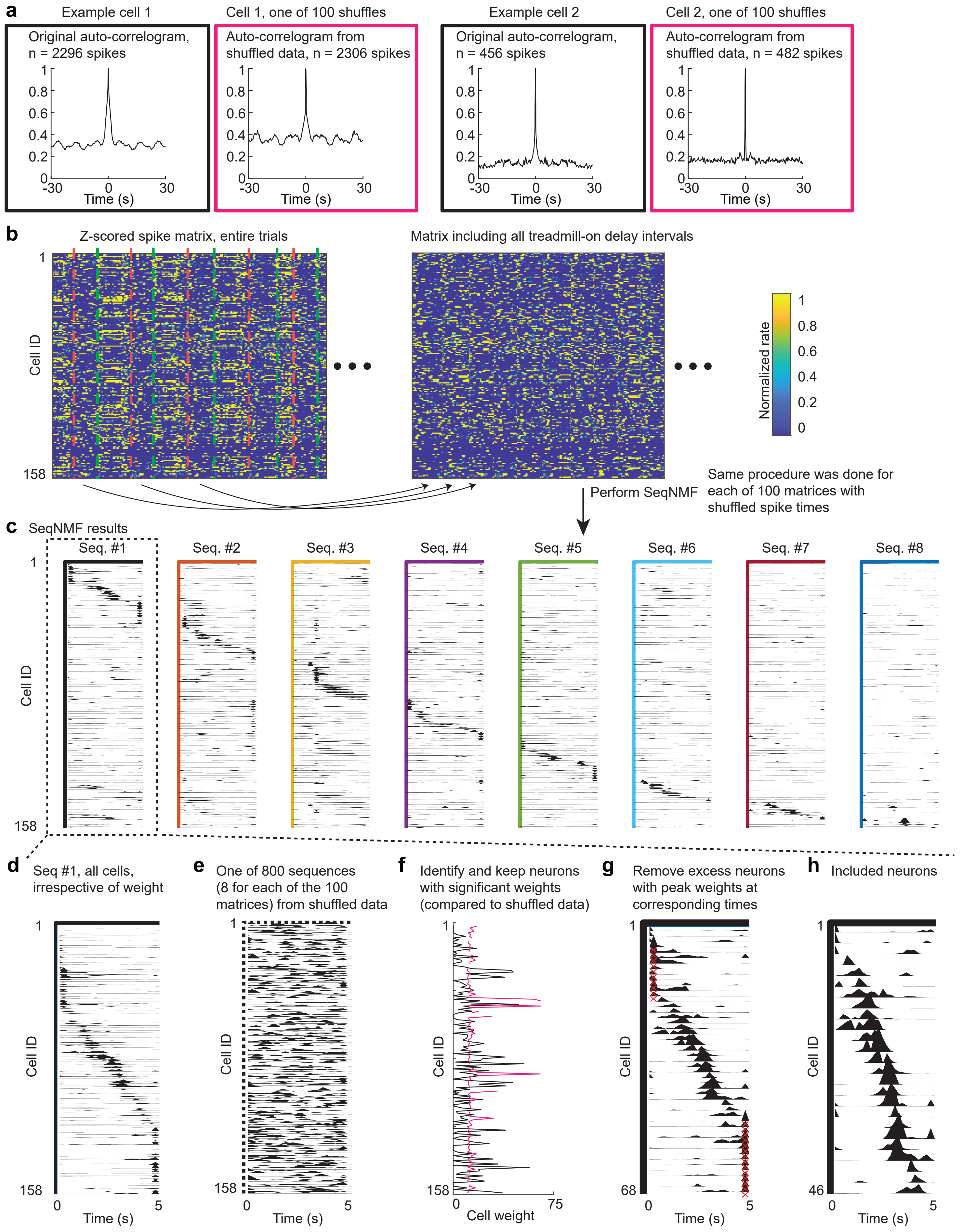

a
Example cell 1
Original auto-correlogram, n = 2296 spikes
Cell 1, one of 100 shuffles
Auto-correlogram from shuffled data, n = 2306 spikes
Example cell 2
Original auto-correlogram, n = 456 spikes
Cell 2, one of 100 shuffles
Auto-correlogram from shuffled data, n = 482 spikes
Time (s)
b
Z-scored spike matrix, entire trials
Matrix including all treadmill-on delay intervals
Cell ID
Normalized rate
Perform SeqNMF
Same procedure was done for each of 100 matrices with shuffled spike times
c
SeqNMF results
Seq. #1
Seq. #2
Seq. #3
Seq. #4
Seq. #5
Seq. #6
Seq. #7
Seq. #8
d
Seq #1, all cells, irrespective of weight
e
One of 800 sequences (8 for each of the 100 matrices) from shuffled data
f
Identify and keep neurons with significant weights (compared to shuffled data)
Cell weight
g
Remove excess neurons with peak weights at corresponding times
h
Included neurons
Time (s)

#### Figure S7, continued

**Figure S7. Sequence detection with SeqNMF and refinement of cell selection. a**, The spike trains of each cell were shuffled using a procedure that preserves the cells' spike rate and temporal autocorrelation. Comparisons of the spike-time autocorrelograms from the original spike train (left of each panel) and of the autocorrelograms from shuffled spike trains (one out of 100 is shown; right of each panel) for two example cells show preserved rate and timing. Note that the shuffling was not merely a time shift, as commonly done, to preclude the possibility that reordered sequences could be detected from shuffled datasets. **b**, Construction of z-scored spike-rate matrices restricted to treadmill-on delay intervals. The same procedure was applied to 100 datasets with shuffled spike trains. **c**, Example result from SeqNMF applied to the original data from a recording session (sequence number = 8). **d**, As shown for the example sequence, all cells were sorted by their peak firing time within the sequence. Note that the analysis assigns a peak time to each cell irrespective of how consistently a cell participated in the sequence. The following step ensured that only cells that consistently participated in the sequence were retained. **e**, Example sequence detected from shuffled data, plotted in the same order as d. For each recording session, 8 sequences were detected from each of the 100 shuffles yielding 800 sequences. **f**, Each cell's weight in the original sequence (black) was compared against the distribution of the same cell's weights from 800 shuffles, and original weights exceeding the 99th percentile of shuffled weights (magenta) were considered significant. For a small number of cells very low firing rates, shuffled weight could not be calculated and are not shown. These neurons were excluded from sequences. **g**, To reduce detection of coincident firing, cells were removed if more than five had peaks in the same time bin. **h**, Final set of cells in the example sequence after applying criteria in f and g.

### Figure S8

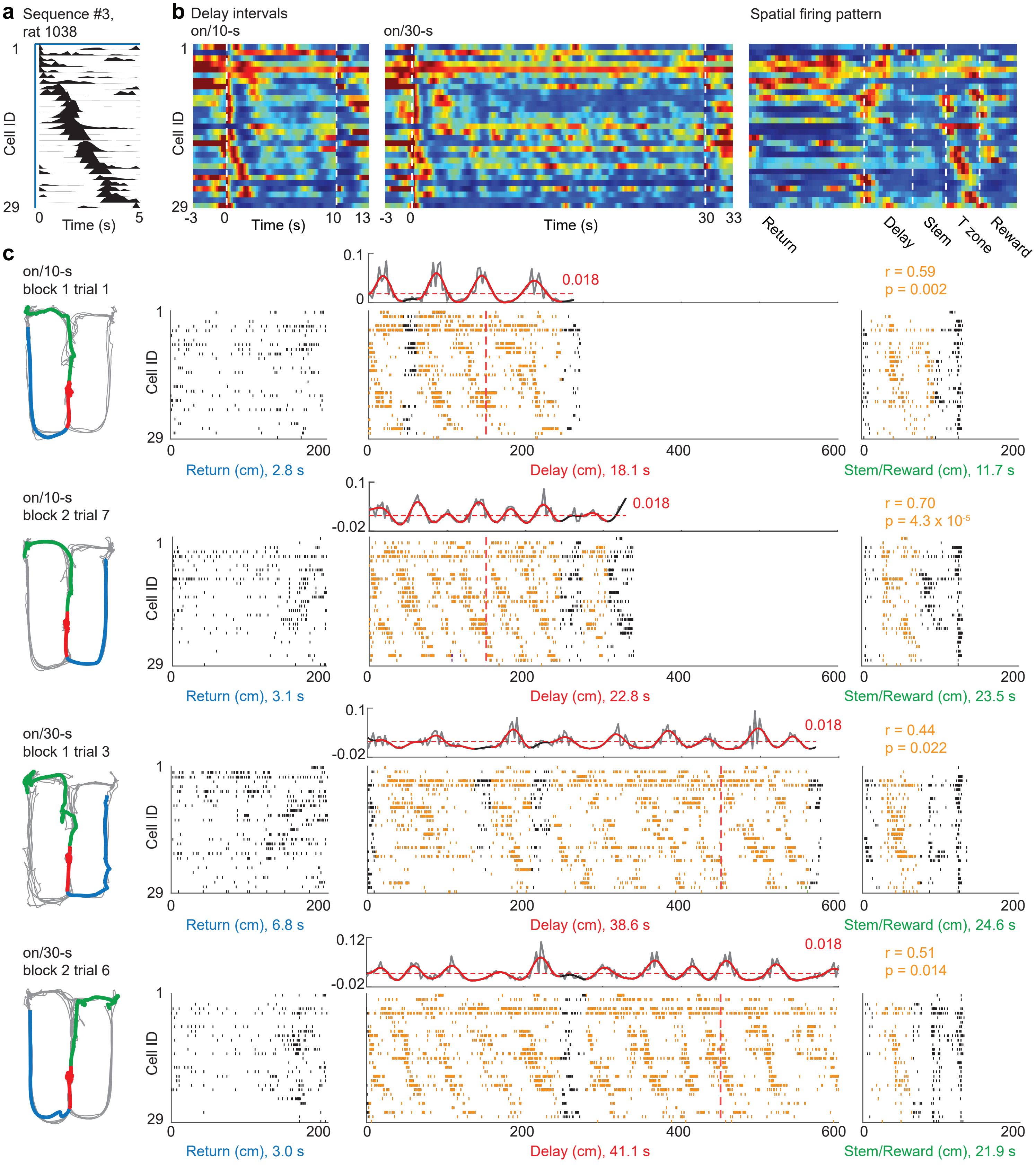


**Figure S8. Example sequence from rat 1138. a**, Neurons participating in the sequence. **b**, Average firing rates of participating neurons during delay intervals and on the maze. Blue to red, 0 Hz to each PN's maximum rate. **c**, Sequence occurrences in example trials. From left to right: animal's trajectory color-coded by maze segment (blue, return; red, delay interval; green, stem to reward), sequence occurrence at return with rank correlation value on top (if sequences are detected), repeating sequences across the delay interval with raw (grey) and smoothed sequence amplitude (red during detected events, otherwise black) on top and spike rasters of participating neurons on bottom, sequence occurrence during the stem-to-reward segment. Cells are ordered as in a, and spikes in/out of the repeating sequence are in orange/black.

#### Figure S9

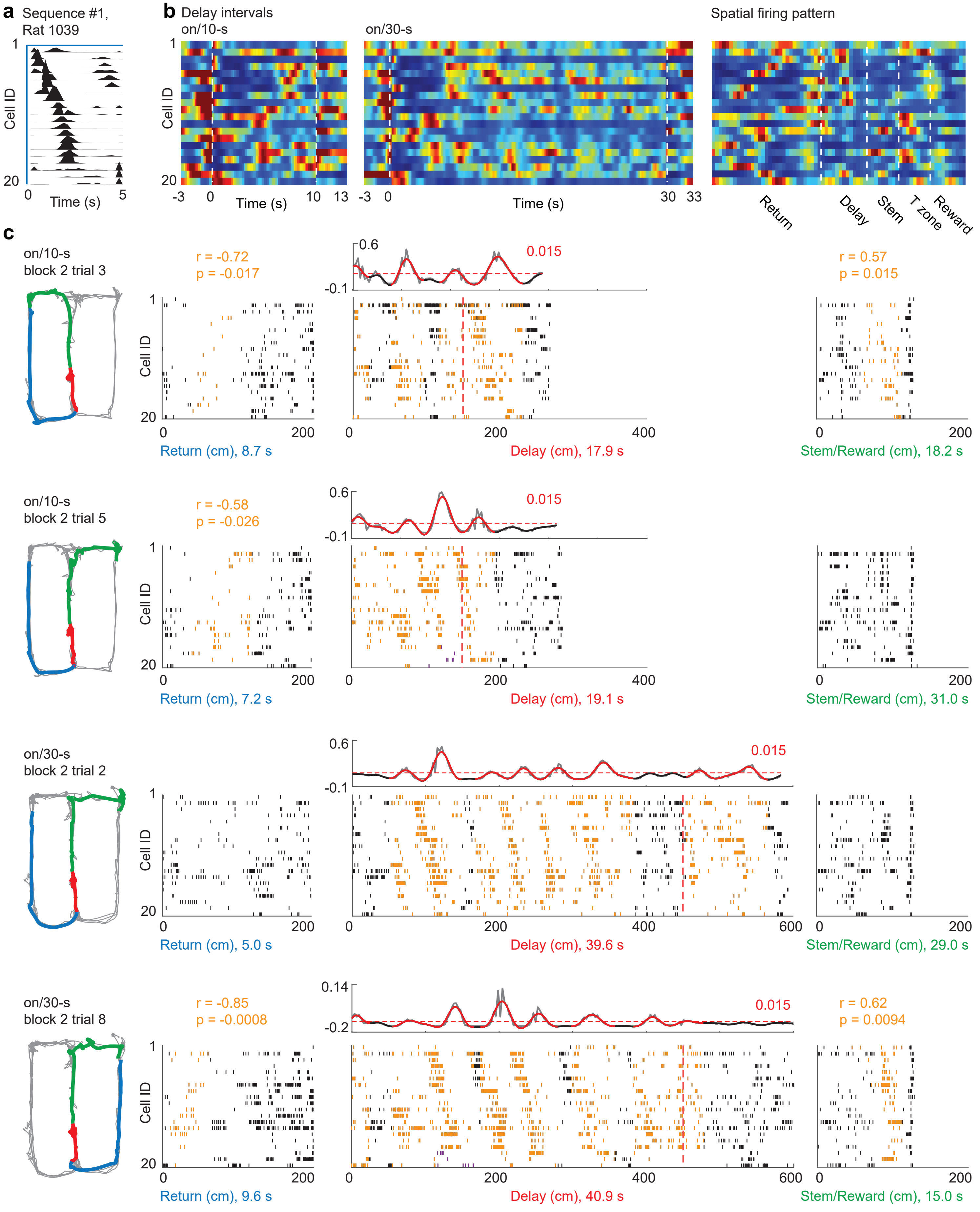


**Figure S9. Example sequence from rat 1139.** Depicted as in Figure S8.

### Figure S10

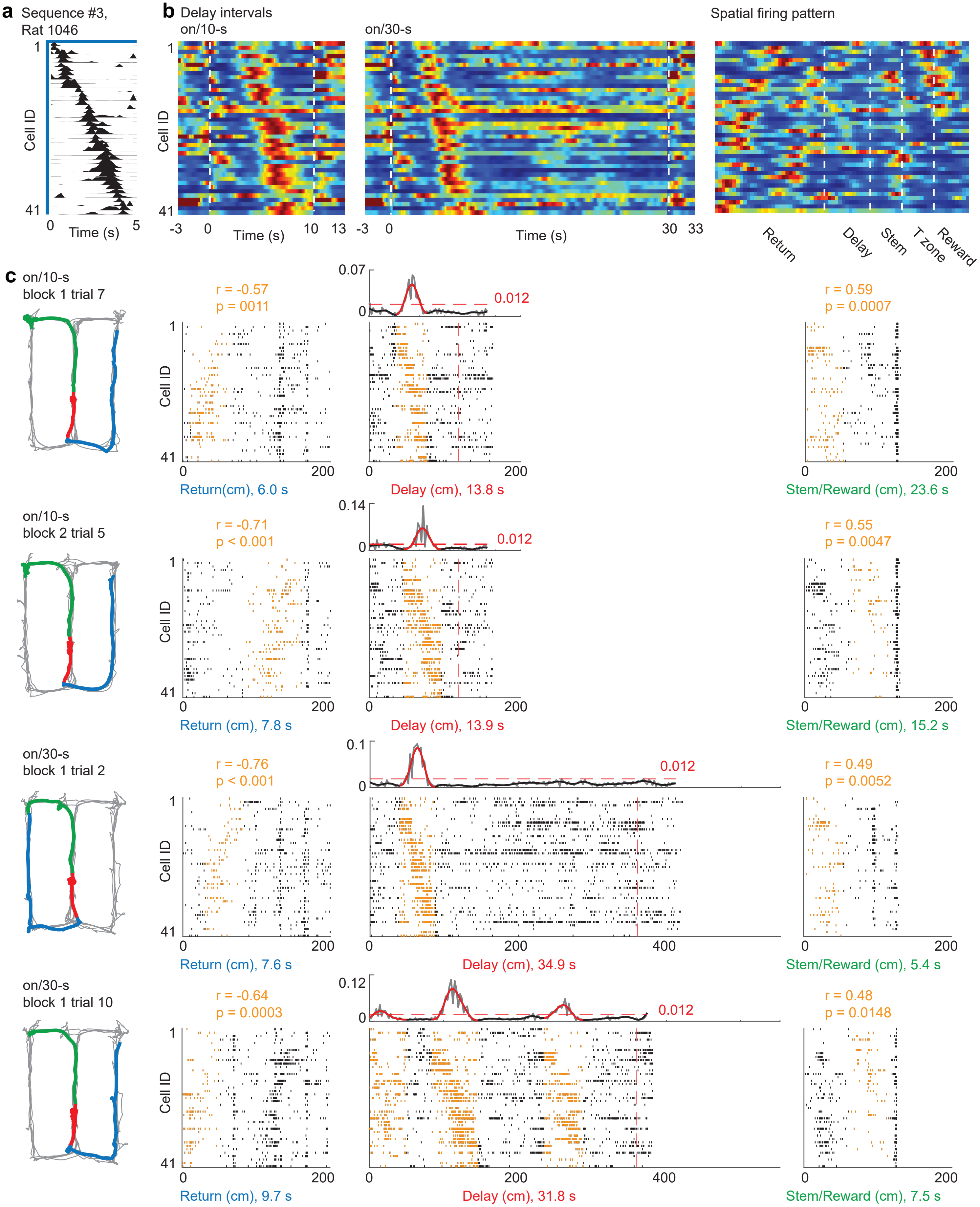


**Figure S10. Example sequence from rat 1146.** Depicted as in Figure S8.

### Figure S11

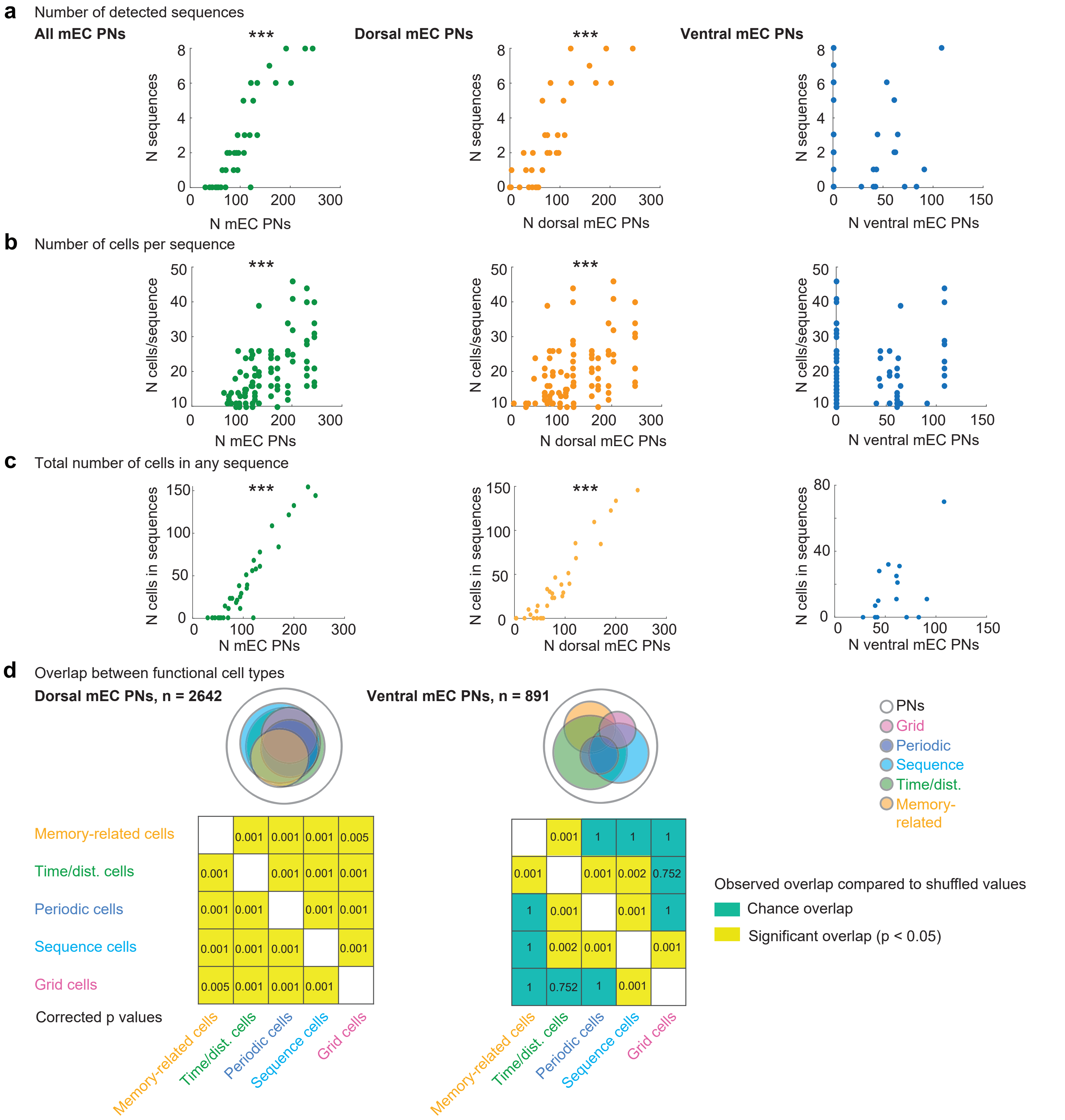


**Figure S11. Relations between the number of recorded cells and sequence metrics. a**, Sequence number increased with the recorded total mEC PNs and dorsal mEC PNs, but not ventral mEC PNs (mEC PNs: $r = 0.897$, $p = 1.6 \times 10^{-12}$; dorsal mEC PNs: $r = 0.859$, $p = 1.5 \times 10^{-10}$; ventral mEC PNs: $r = -0.141$, $p = 0.433$; $n = 33$, Spearman's correlation). **b**, Detected number of cells in each sequence increased with the recorded total mEC PNs and dorsal mEC PNs, but not ventral mEC PNs (mEC PNs: $r = 0.612$, $p = 7.4 \times 10^{-11}$; dorsal mEC PNs: $r = 0.491$, $p = 5.7 \times 10^{-7}$; ventral mEC PNs: $r = -0.004$, $p = 0.968$; $n = 93$, Spearman's correlation). **c**, Detected number of cells in any sequence increased with the recorded total mEC PNs and dorsal mEC PNs, but not ventral mEC PNs (mEC PNs: $r = 0.914$, $p = 1.0 \times 10^{-13}$; dorsal mEC PNs: $r = 0.919$, $p = 3.1 \times 10^{-13}$; ventral mEC PNs: $r = 0.402$, $p = 0.137$; $n = 33$, Spearman's correlation). *** $p < 0.001$. **d**, Overlap between functional cell types. Substantial overlap among all identified cell types was observed in dorsal mEC (adjusted permutation test, $p < 0.05$ for all comparisons), while the small proportion of some cell types (e.g., grid cells, periodic cells) in ventral mEC resulted in more limited overlap (see figure for statistics).

#### Figure S12

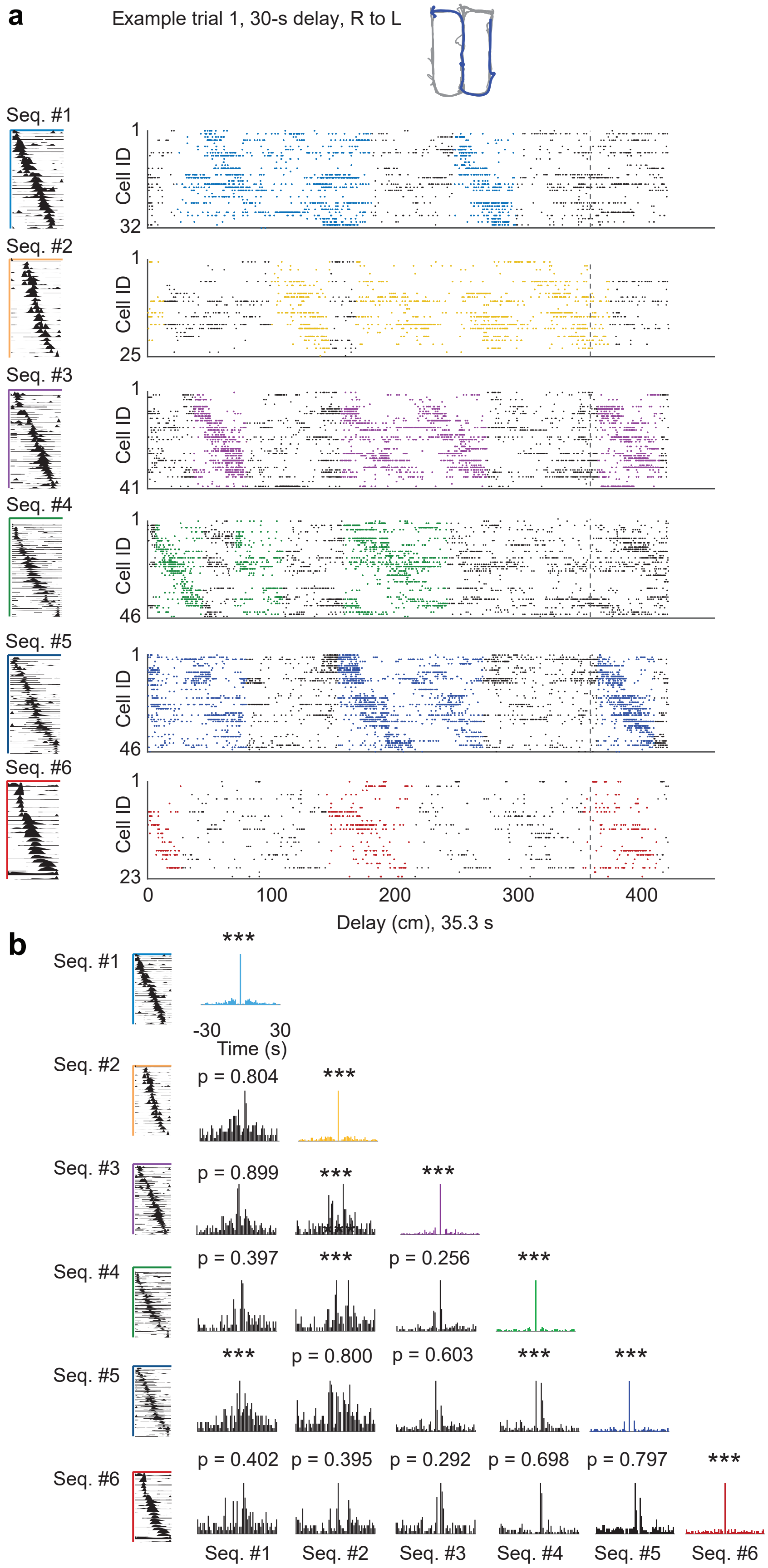


**Figure S12. Multiple distinct mEC sequences played out throughout delay intervals. a**, Two example trials from a recording session with six distinct sequences throughout the delay interval. For each sequence, the left panel shows the SeqNMF-derived weight profile across neurons, and the panels to the right show spike rasters of participating neurons, ordered as shown to the left. **b**, Autocorrelations reveals robust repetition for individual sequences, whereas cross-correlations show only limited temporal coupling between different sequences. *** $p < 0.001$.

#### Figure S13

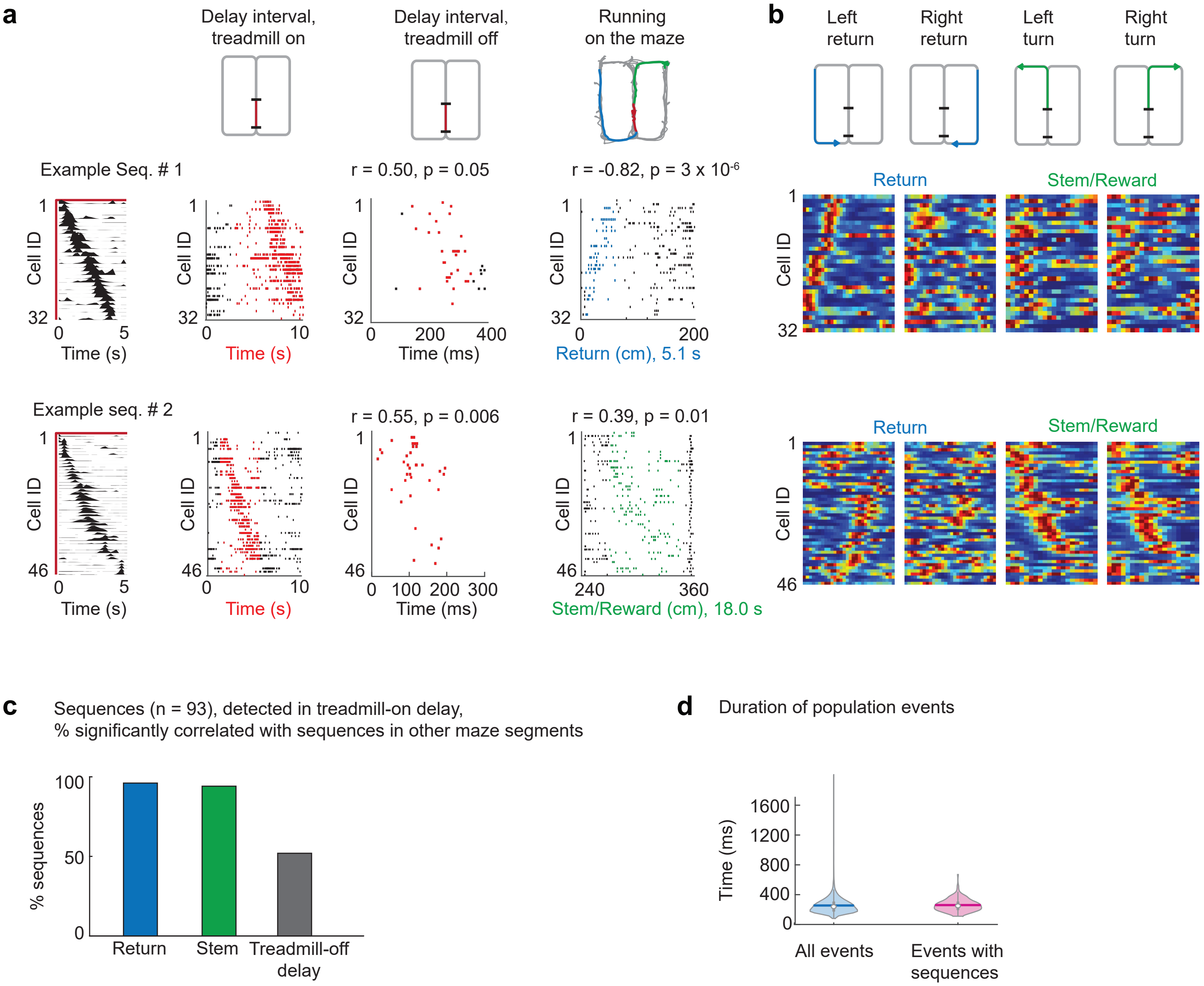


**Figure S13. Sequences that were identified during treadmill-on delays also played out in other task phases. a**, From left to right, sequence template, sequence during the treadmill-on delay of a single trial, sequential activation of the same cells during the treadmill-off delay interval, sequential activation of the same cells on the return arm in reverse order (top, return) or after exiting the delay zone in forward order (bottom, stem to reward). Top and bottom rows show two different sequences from the same recording session. **b**, Spatial maps for the cells in the sequences shown in a. Each line is a cell, and color maps are from 0 Hz (blue) to maximum firing rate (red). Maps are generated by averaging over all trials in a condition (left return, right return, left turn, right turn) and include the maze segment corresponding to the colored arrow in the schematics on top (left two panels, return; right two panels, stem to reward). **c**, Percent sequences (detected in treadmill-on delay intervals) that are significantly correlated with sequential activity in other phases (return arms, stem/T zone or treadmill-off delay intervals; n = 93 sequences). **d**, Length of all population events (i.e., brief periods of high principal cell activity) and of population events in which sequential activity patterns could be detected (mean ± SEM: 254 ms ± 1.1 and 260 ms ± 6.2). Violin plot: width, horizontal line, open circle, thick vertical line and thin vertical line; distribution, mean, median, 25th/75th quartile and maximum/minimum except outliers.

#### Figure S14

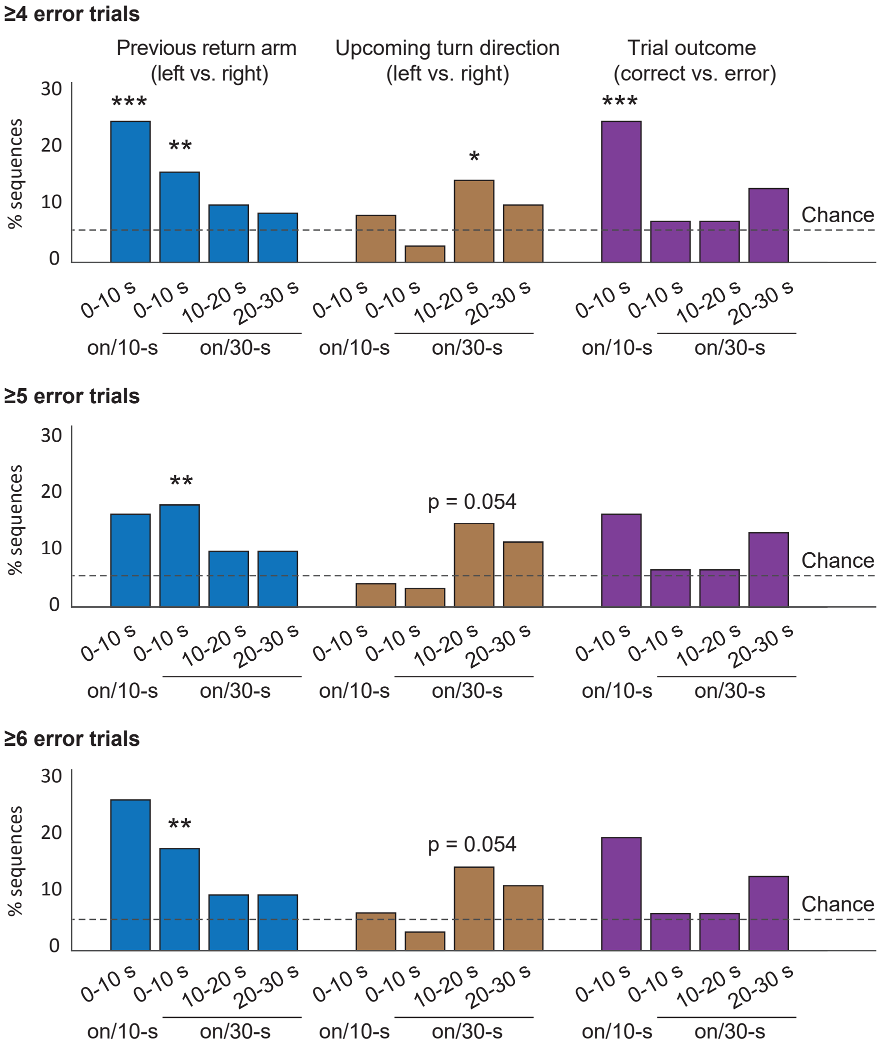


**Figure S14. Coding for memory-related information is robust across different error thresholds.** Treadmill-on delay sequences can preferentially be active depending on the preceding return arm [left (L) vs. right (R)], the upcoming turn (L vs. R) or the trial outcome (correct vs. error). These effects were observed consistently with increasingly stringent minimum errors in a delay condition.

Minimum of 4 errors, return L vs. R: on/10-s: 23.53%, p = 0.00007, n = 51; on/30-s: 0-10 s, 15.07%, p = 0.010; 10-20 s, 9.59%, p = 0.67; 20-30 s, 8.22%, p = 0.91, n = 73 sequences; upcoming L vs. R turn: on/10-s: 7.84%, p = 1, n = 51 sequences; on/30-s: 0-10 s, 2.74%, p = 0.59; 10-20 s, 13.70%, p = 0.03; 20-30 s, 9.59%, p = 0.58, n = 73 sequences; outcome correct vs. error: on/10-s: 23.53%, p = 0.00007, n = 51 sequences; on/30-s: 0-10 s, 6.85%, p = 0.42; 10-20 s, 6.85%, p = 0.83; 20-30 s, 12.33%, p = 0.08, n = 73 sequences; binomial test compared to chance proportion of 5%, p values corrected.

Minimum of 5 errors, return L vs. R: on/10-s: 15.39%, p = 0.35, n = 26; on/30-s: 0-10 s, 16.92%, p = 0.004; 10-20 s, 9.23%, p = 0.71; 20-30 s, 9.23%, p = 0.71, n = 65 sequences; upcoming L or R turn: on/10-s: 3.85%, p = 1, n = 26 sequences; on/30-s: 0-10 s, 3.08%, p = 1; 10-20 s, 13.85%, p = 0.054; 20-30 s, 10.77%, p = 0.30, n = 65 sequences; outcome correct vs. error: on/10-s: 15.39%, p = 0.31, n = 26 sequences; on/30-s: 0-10 s, 6.15%, p = 1; 10-20 s, 6.15%, p = 1; 20-30 s, 12.31%, p = 0.14, n = 65 sequences; binomial test compared to chance proportion of 5%, p values corrected.

Minimum 6 errors, return L vs. R: on/10-s: 25.00%, p = 0.07, n = 16 sequences; on/30-s: 0-10 s, 16.92%, p = 0.004; 10-20 s, 9.23%, p = 0.71; 20-30 s, 9.23%, p = 0.71, n = 65 sequences; upcoming L vs. R turn: on/10-s: 6.25%, p = 1, n = 16 sequences; on/30-s: 0-10 s, 3.08%, p = 1; 10-20 s, 13.85%, p = 0.054; 20-30 s, 10.77%, p = 0.30, n = 65 sequences; outcome correct vs. error: on/10-s: 18.75%, p = 0.34, n = 16; on/30-s: 0-10 s, 6.15%, p = 1; 10-20 s, 6.15%, p = 1; 20-30 s, 12.31%, p = 0.14, n = 65 sequences; binomial test compared to chance proportion of 5%, p values corrected. * p < 0.05, ** p < 0.01, *** p < 0.001.

**Table S1. Numbers of recorded dorsal and ventral mEC principal neurons (PNs) per session**

| Session | Rat | Recording ID | Postsurg. day | Number of recorded mEC PNs | | |
|---|---|---|---|---|---|---|
| | | | | dorsal | ventral | total |
| 1 | 1140 | 20240108 | 4 | 38 | 83 | 121 |
| 2 | 1140 | 20240109 | 5 | 28 | 61 | 89 |
| 3 | 1140 | 20240110 | 6 | 20 | 42 | 62 |
| 4 | 1140 | 20240111 | 7 | 3 | 28 | 31 |
| 5 | 1140 | 20240112 | 8 | 0 | 40 | 40 |
| 6 | 1137 | 20240205 | 4 | 4 | 91 | 95 |
| 7 | 1137 | 20240208 | 7 | 0 | 71 | 71 |
| 8 | 1139 | 20240304 | 5 | 95 | 0 | 95 |
| 9 | 1139 | 20240305 | 6 | 75 | 0 | 75 |
| 10 | 1139 | 20240306 | 7 | 65 | 0 | 65 |
| 11 | 1139 | 20240307 | 8 | 54 | 0 | 54 |
| 12 | 1138 | 20240401 | 4 | 97 | 0 | 97 |
| 13 | 1138 | 20240402 | 5 | 171 | 0 | 171 |
| 14 | 1138 | 20240403 | 6 | 107 | 0 | 107 |
| 15 | 1138 | 20240404 | 7 | 122 | 0 | 122 |
| 16 | 1138 | 20240405 | 8 | 109 | 0 | 109 |
| 17 | 1147 | 20240527 | 4 | 45 | 0 | 45 |
| 18 | 1147 | 20240528 | 5 | 52 | 0 | 52 |
| 19 | 1147 | 20240529 | 6 | 57 | 0 | 57 |
| 20 | 1148 | 20240624 | 4 | 65 | 61 | 126 |
| 21 | 1148 | 20240625 | 5 | 32 | 40 | 72 |
| 22 | 1146 | 20240708 | 4 | 244 | 0 | 244 |
| 23 | 1146 | 20240710 | 6 | 191 | 0 | 191 |
| 24 | 1146 | 20240711 | 7 | 201 | 0 | 201 |
| 25 | 1146 | 20240712 | 8 | 158 | 0 | 158 |
| 26 | 1151 | 20241028 | 4 | 44 | 43 | 87 |
| 27 | 1151 | 20241029 | 5 | 70 | 64 | 134 |
| 28 | 1151 | 20241030 | 6 | 46 | 62 | 108 |
| 29 | 1151 | 20241031 | 7 | 81 | 53 | 134 |
| 30 | 1150 | 20241119 | 5 | 121 | 108 | 229 |
| 31 | 1150 | 20241120 | 6 | 75 | 44 | 119 |
| 32 | 1150 | 20241122 | 7 | 79 | 0 | 79 |
| 33 | 1150 | 20241123 | 8 | 93 | 0 | 93 |

**Table S2. Proportion of cells with delay activity depending on previous return arm and/or upcoming turn direction.**

| | | | | 30-s delay intervals | | | | | |
|---|---|---|---|---|---|---|---|---|---|
| N cells/sessions | N rats | Condition | | 0-5s % | 5-10s % | 10-15s % | 15-20s % | 20-25s % | 25-30s % |
| Trajectory dependent (L-R vs. R-L) | | | | | | | | | |
| 1758/18[1] | 9 | on/30-s | Mean | 10.81 | 5.06 | 5.63 | 4.61 | 4.72 | 4.44 |
| | | | p[2] | $3.9 \times 10^{-22}$ | 0.87 | 1 | 1 | 1 | 1 |
| 2834/27[1] | 9 | off/30-s | Mean | 11.58 | 8.21 | 6.48 | 5.21 | 5.86 | 6.11 |
| | | | p[2] | $8.8 \times 10^{-42}$ | $1.2 \times 10^{-11}$ | 0.0059 | 1 | 0.28 | 0.070 |
| Previous return arm (L vs. R) | | | | | | | | | |
| 1775/15[3] | 6 | on/30-s | Mean | 12.34 | 5.80 | 4.68 | 4.68 | 5.18 | 3.83 |
| | | | p[2] | $4.8 \times 10^{-34}$ | 0.64 | 1 | 1 | 1 | 0.13 |
| 769/6[3] | 5 | off/30-s | Mean | 15.21 | 11.44 | 6.89 | 8.45 | 5.07 | 6.89 |
| | | | p[2] | $8.1 \times 10^{-17}$ | $1.1 \times 10^{-11}$ | 0.16 | 0.56 | 0.93 | 0.14 |
| Upcoming turn selective (L vs. R) | | | | | | | | | |
| 1775/15[3] | 6 | on/30-s | Mean | 4.00 | 4.11 | 4.45 | 5.75 | 6.25 | 5.97 |
| | | | p[2] | 0.45 | 0.55 | 0.33 | 0.47 | 0.19 | 0.45 |
| 769/6[3] | 5 | off/30-s | Mean | 11.57 | 6.11 | 6.24 | 7.02 | 6.24 | 6.63 |
| | | | p[2] | $5.9 \times 10^{-12}$ | 0.32 | 0.58 | 0.14 | 0.46 | 0.42 |

[1] with ≤5 error trials in 20 trials, [2]compared to chance level (5%), Holm-Bonferroni corrected, [3] with ≥6 error trials in 20 trials,
